# Selective GSK3α Inhibition Promotes Self-Renewal Across Different Stem Cell States

**DOI:** 10.1101/2025.05.16.653860

**Authors:** Duo Wang, Xiukun Wang, Shuling Wang, Kai-Xuan Shi, Safia Malki, Yanpui Chan, Joshua Feng, Jiaqi Tang, Xi Chen, Daniel McKim, Chao Zhang, Guang Hu, Qi-Long Ying

**Author notes:** These authors contributed equally (Duo Wang, Xiukun Wang).

## Abstract

Pan-GSK3α/β inhibition promotes stem cell self-renewal through activation of WNT/β-catenin signaling, but its broad effects complicate the precise control of stem cell states. Here, we show that selective inhibition of GSK3α with BRD0705 supports the long-term self-renewal of mouse embryonic stem cells (ESCs), epiblast stem cells (EpiSCs), and neural stem cells (NSCs), independent of β-catenin signaling. When combined with the tankyrase inhibitor IWR1, BRD0705 broadly supports the maintenance of diverse pluripotent stem cell states, including ESCs, EpiSCs, and formative pluripotent stem cells. This BRD0705/IWR1 cocktail enables stable co-culture of naive ESCs and primed EpiSCs while preserving their distinct molecular and functional identities. Single-cell transcriptomics, epigenomic profiling, and functional assays confirm sustained lineage-specific features across stem cell types. These findings demonstrate that selective GSK3α inhibition enhances stemness by buffering against differentiation cues and promoting intrinsic self-renewal capacity. This work identifies GSK3α as a key regulator of self-renewal across distinct stem cell states and establishes a versatile culture system with broad applications.

**In Brief:** Wang et al. demonstrate that selective GSK3α inhibition with BRD0705 supports self-renewal of pluripotent and neural stem cells. Combined with IWR1, it enables long-term co-culture of naive and primed stem cells while preserving their distinct molecular and functional identities.

**Graphical abstract:** 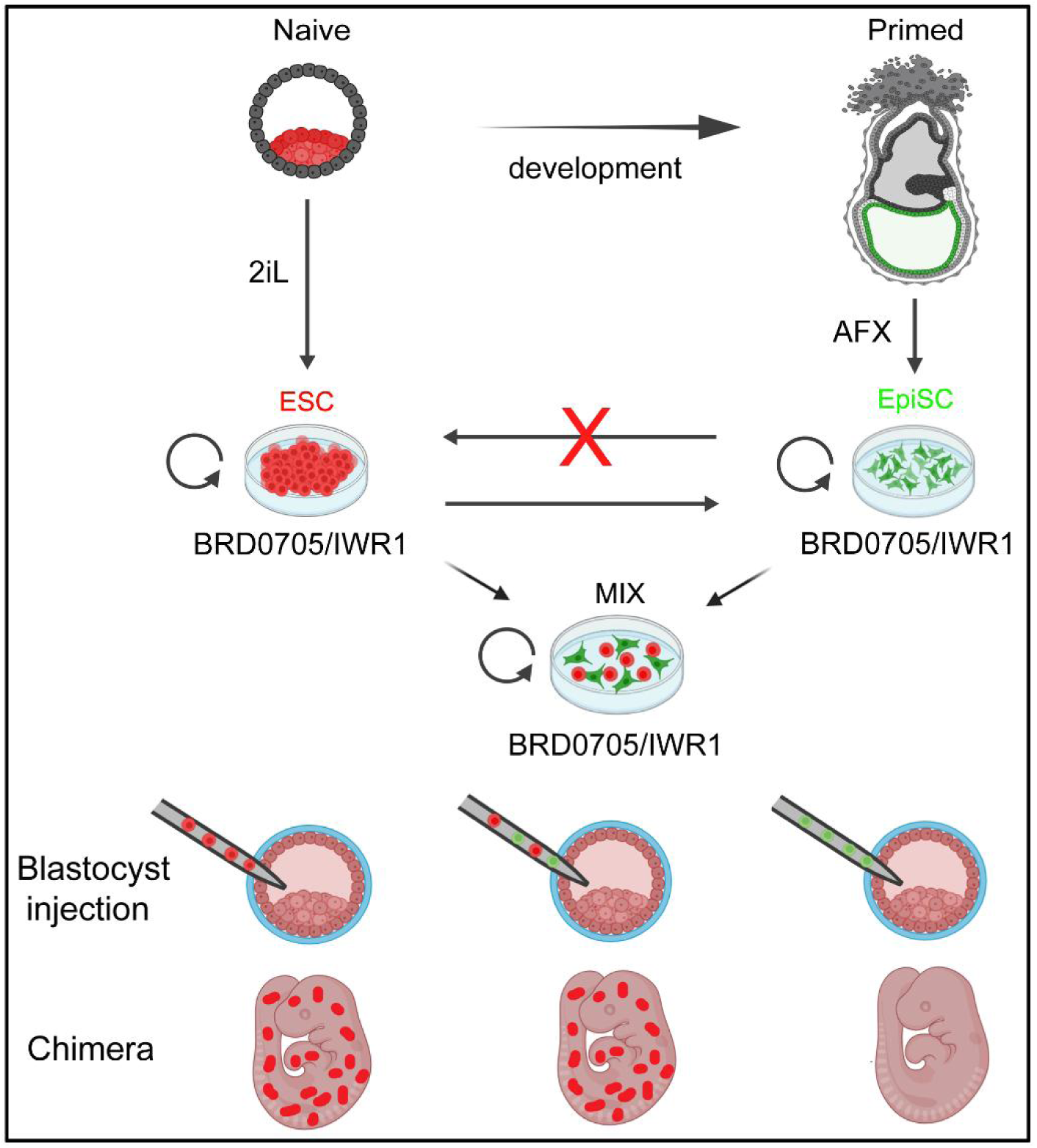

**Highlights:**

- GSK3α inhibition by BRD0705 promotes self-renewal of ESCs, EpiSCs, and NSCs
- BRD0705/IWR1 enables long-term co-culture of ESCs and EpiSCs
- Co-cultured ESCs and EpiSCs retain distinct naive or primed identities
- BRD0705 preserves stem cell states independently of β-catenin signaling

## Introduction

Pluripotency is defined as a cell’s ability to differentiate into all three germ layers: endoderm, mesoderm, and ectoderm.^1, 2^ Mouse embryonic stem cells (ESCs) and epiblast stem cells (EpiSCs) represent the two most extensively studied types of pluripotent stem cells (PSCs).^3–5^ ESCs are derived from the pre-implantation epiblast and exist in a naive state, characterized by their capacity to contribute extensively to chimeric embryos and transmit through the germline upon blastocyst injection.^6–10^ In contrast, EpiSCs are isolated from the post-implantation epiblast and represent a primed state of pluripotency.^11, 12^ While EpiSCs can differentiate into derivatives of all three germ layers in vitro, they exhibit limited developmental potential in vivo and fail to generate high-contribution chimeras.^13–15^ Consequently, EpiSCs are classified as being in a primed pluripotent state.^7^ *In vitro*, naive ESCs readily acquire features of the primed state under appropriate conditions, mirroring the in vivo transition from pre- to post-implantation epiblast.^16^ However, reversion of primed EpiSCs to the naive state typically requires the forced expression of transcription factors, suggesting that distinct molecular mechanisms govern self-renewal in these two pluripotent states.^16, 17^

Indeed, the maintenance of naive and primed PSCs depends on fundamentally different, and often antagonistic, signaling environments.^7, 9, 18, 19^ Naive ESCs are propagated under conditions that include the MEK inhibitor PD0325901 (PD03) and the GSK3 inhibitor 99021 (CHIR), a combination known as 2i,^8, 20^ which blocks differentiation-inducing FGF/MEK signaling^6, 9^ and activates WNT/β-catenin signaling^9, 21, 22^, respectively. In contrast, primed EpiSCs require culture in medium containing Activin A, bFGF, and the WNT inhibitor XAV939 (AFX), creating an opposing signaling environment.^16, 23, 24^ This divergence not only reflects distinct developmental states but also presents a practical challenge: ESCs readily convert to EpiSCs under AFX conditions, but EpiSCs cannot be maintained in 2i or revert back to naive ESCs without reprogramming interventions.^16, 17^ Whether naive and primed PSCs share any common molecular pathways that support pluripotency remains unknown.

While primed PSCs have been established from a wide range of mammalian species,^25–28^ the derivation of true naive ESCs, defined by robust chimera formation and germline transmission, has thus far been successful only in mice and rats.^9, 29, 30^ Notably, the 2i condition that enables long-term propagation of mouse and rat ESCs fails to support ESC derivation from most other species, including humans.^9, 30–33^ This limitation suggests that alternative or additional signaling inputs may be required to sustain naive pluripotency in a broader range of mammals.

Identifying core self-renewal mechanisms shared across naive and primed PSCs could provide a unifying framework for pluripotent stem cell maintenance and offer critical insight into the molecular requirements for deriving chimera-competent ESCs from other species. The development of such universal culture conditions would have far-reaching implications for regenerative medicine, species conservation, and developmental biology.

GSK3 is a highly conserved serine/threonine kinase family central to many cellular signaling pathways and essential for diverse cellular functions.^34–39^ In mammals, two paralogs exist: GSK3α and GSK3β.^40^ In naive ESCs, selective inhibition of GSK3β suppresses β-catenin phosphorylation, thereby maintaining ESC pluripotency, similar to the effect observed with CHIR;^22^ however, the role of GSK3α in regulating ESCs fate remains largely unexplored.

In this study, we screened small-molecule inhibitors to identify compounds capable of supporting the self-renewal of both naive ESCs and primed EpiSCs. We identified BRD0705, a selective GSK3α inhibitor,^41^ as a key factor that promotes the propagation of both cell types. When combined with the tankyrase inhibitor IWR1, BRD0705 enables the long-term culture of naive ESCs, primed EpiSCs, and formative pluripotent cells while preserving their lineage-specific molecular and functional characteristics. Notably, ESCs cultured in BRD0705/IWR1 retain full developmental potential, contributing to high-level chimeras and germline transmission. Our findings reveal a previously unrecognized role for GSK3α in PSC self-renewal and establish BRD0705/IWR1 as a broadly applicable culture condition for maintaining diverse pluripotent states.

## Results

### BRD0705 promotes self-renewal of both ESCs and EpiSCs

We screened a small-molecule library (Table S1) using ESCs to identify novel compounds that promote ESC self-renewal and to gain insights into the underlying regulatory mechanisms. The library included compounds targeting key regulators of cell proliferation and stemness, including protein tyrosine kinases and components of the WNT, MAPK, PI3K/AKT/mTOR, TGFβ/Smad, JAK/STAT, and NF-κB signaling pathways. In the initial screen, 47 compounds enhanced ESC colony formation (Figure S1A). These hits were further evaluated by alkaline phosphatase (AP) staining, which identified BRD0705, a selective GSK3α inhibitor,^41^ as the most effective molecule in sustaining ESC self-renewal in the absence of 2i/LIF (2iL) (Figures 1A and 1B). After extended passaging, BRD0705 was the only compound that consistently maintained ESC self-renewal over time (Figures 1C, S2A, and S2B).

**Figure 1.**
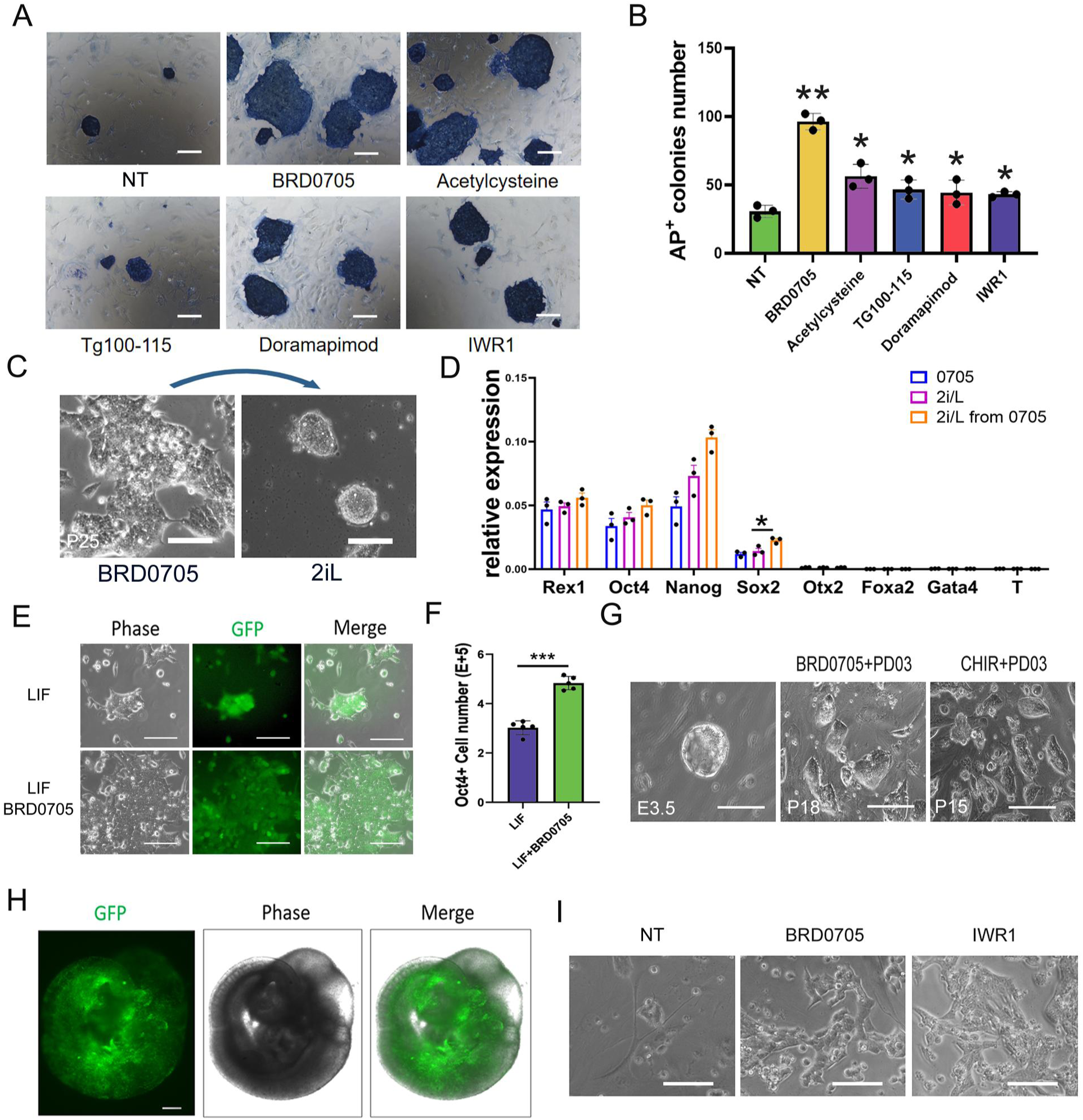
BRD0705 Effectively Supports the Self-renewal of Both ESCs and EpiSCs. (A) Alkaline phosphatase (AP) staining of ESC colonies formed under treatment with various compounds from the chemical inhibitor library. Cells were cultured in DMEM/FBS medium for 4 days. Scale bar, 100 μm. NT, no treatment control. (B) Quantification of AP+ colonies. Data represent mean±SEM. from triplicate experiments. *, p<0.05, indicating a statistically significant difference in the number of AP+ clones compared to the control under different inhibitor treatments. (C) Left panel: a representative phase-contrast image showing the morphology of ESCs at passage 25 cultured in DMEM/10%FBS under BRD0705 treatment. Right panel: A representative phase-contrast image showing the morphology of ESCs cultured in 2iL conditions for three passages after being maintained in BRD0705 for 25 passages. Scale bar: 100 μm. (D) qRT-PCR analysis of pluripotency and germ layer-specific gene expression in ESCs maintained in 2iL, BRD0705, or in cells transitioned from BRD0705 to 2iL, normalized to GAPDH. Error bars are SEM from technical triplicates. *, p<0.05. (E) Representative phase-contrast and fluorescence images of Oct4-GFP ESCs cultured in LIF alone or with BRD0705.The basal cell culture medium was DMEM/10%FBS. Scale bar, 100 μm. (F) Quantification of Oct4-GFP+ colony ratio from the experiments shown in (E). Data are presented as mean ± SEM from three biological replicates. *, p<0.05. **, p<0.01. ***, p<0.001. (G) Phase-contrast images of mouse embryo blastocysts (E3.5), passage 18 ESCs derived from blastocysts under BRD0705+PD03 conditions, and ESCs cultured under CHIR+PD03 conditions. Scale bar: 200 μm. (H) Bright-field and fluorescent images of E10.5 mouse embryos generated after blastocyst injection of BRD0705+PD03 cultured ESCs with ubiquitous GFP. Scale bars, 350 μm. (I) Representative phase-contrast images of EpiSCs treated with BRD0705 or IWR1 from the compound library on day 3. Scale bar 100μm.

Surprisingly, 129P2/Ola strain mESCs cultured with BRD0705 alone remained undifferentiated for over 25 passages (Figure 1C, left panel). These cells also transitioned directly to the 2iL condition, suggesting retention of naive pluripotency (Figure 1C, right panel). Similar results were observed in ESCs derived from a different strain (B6D2F1), which sustained long-term self-renewal in BRD0705 in both serum-containing and serum-free media (Figure S2C).

Quantitative RT-PCR analysis confirmed that ESCs maintained in BRD0705 expressed pluripotency genes (*Rex1, Oct4*, *Nanog, Sox2*) at levels comparable to those in 2iL controls (Figure 1D). ESCs transferred from BRD0705 to 2iL retained this expression pattern (Figure 1D), further supporting the ability of BRD0705 to maintain pluripotency.

To further investigate the role of BRD0705 in maintaining pluripotency, we utilized Oct4-GIP ESCs, in which GFP expression is driven by the Oct4 promoter.^42^ Under LIF/serum conditions, treatment with BRD0705 significantly enhanced the expansion of Oct4-GFP+ cells (Figures 1E and 1F). Furthermore, BRD0705 effectively substituted for CHIR in N2B27+PD03 medium, enabling the derivation and stable expansion of B6D2F1 ESC lines (Figure 1G). When ESCs with ubiquitous GFP expression, cultured in BRD0705/PD03, were injected into wild-type blastocysts, they exhibited robust chimeric competence. Notably, 57.1% of embryos injected with these cells developed into chimeric embryos at E10.5 (Figures 1H and S2D).

Using the same pool of small molecules (Table S1), we conducted a parallel screen on mouse EpiSC cultured in DMEM/10% FBS. This screen identified 21 compounds that promoted EpiSC self-renewal (Figure S1B). Remarkably, despite screening EpiSCs rather than ESCs, BRD0705 again emerged as one of the top candidates (Figures 1I and S1B). Taken together, these findings demonstrate that BRD0705 support the self-renewal of both ESCs and EpiSCs.

Although BRD0705 enhanced EpiSC self-renewal, EpiSCs cultured in DMEM/10% FBS with BRD0705 alone gradually differentiated and could not be maintained beyond five passages (Figure S2E). To further optimize conditions for EpiSC self-renewal, we tested BRD0705 in combination with additional small molecules. In our screen, the tankyrase inhibitor IWR1 was identified as a promoter of EpiSC self-renewal (Figures 1I and S1B). We therefore examined the combined effect of BRD0705 and IWR1 on EpiSCs. This combination supported robust, long-term self-renewal of EpiSCs (Figure 2A). Interestingly, IWR1 also modestly enhanced ESC self-renewal (Figures 1A, 1B, and S1A), prompting us to evaluate the BRD0705/IWR1 combination in ESCs. Unexpectedly, this combination not only maintained EpiSCs but also supported the long-term expansion of ESCs (Figure 2A).

**Figure 2.**
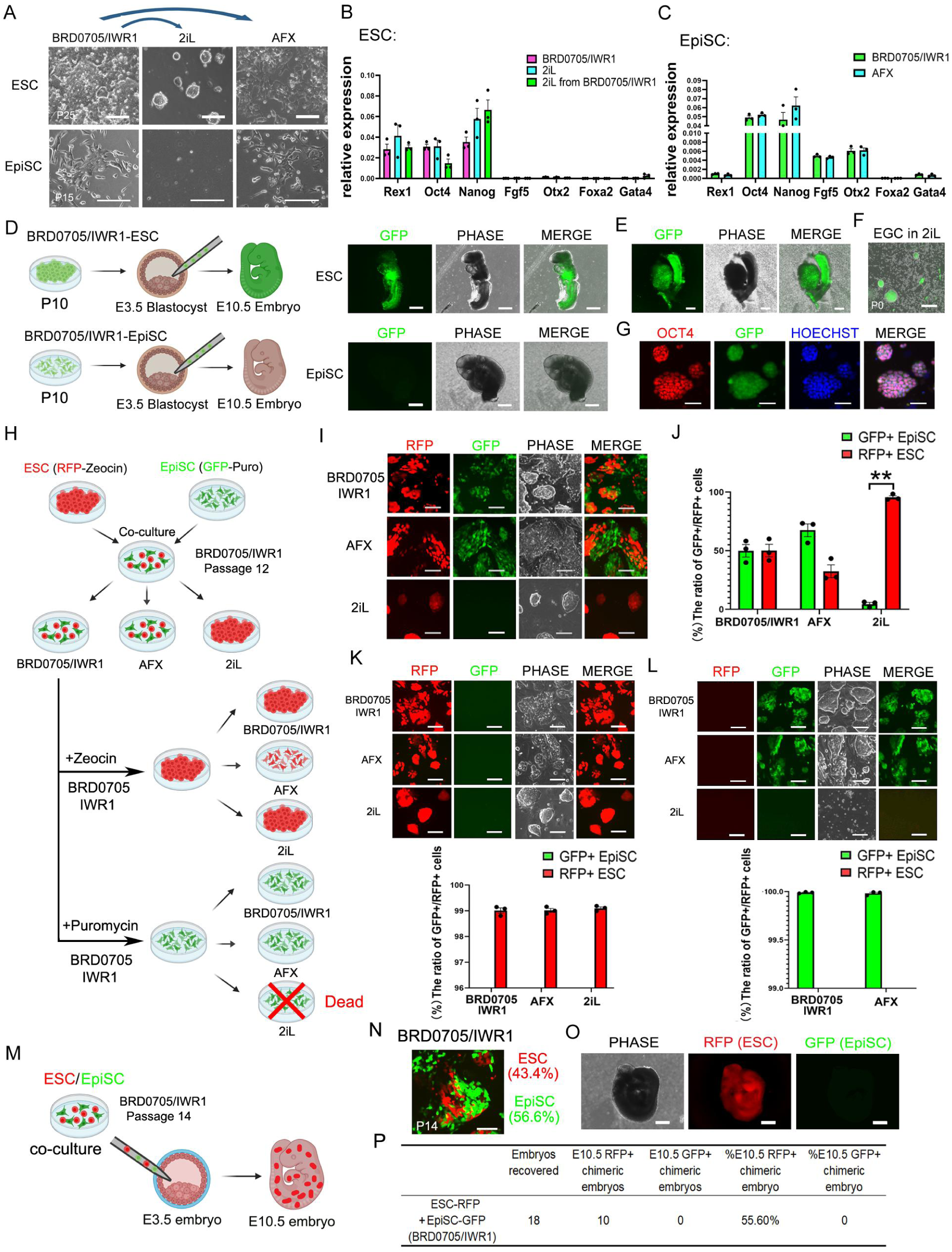
Characterization of ESCs and EpiSCs Cultured under BRD0705/IWR1. (A) Top panel: Phase-contrast images showing ESCs cultured in BRD0705/IWR1 for 25 passages, then transitioned to 2iL or ActivinA/bFGF/XAV condition for 3 passages (scale bar: 100 μm). Bottom panel: phase-contrast images showing EpiSCs cultured 15 passages in BRD0705/IWR1, then transitioned to 2iL or ActivinA/bFGF/XAV conditions for 3 passages (scale bar: 200 μm). (B) qRT-PCR analysis of marker gene expression in ESCs cultured in BRD0705/IWR1 for 15 passages, 2iL, and 2iL after transition from BRD0705/IWR1-cultured ESCs, normalized to GAPDH. Data represent mean ± SEM, n=3.qRT-PCR analysis of marker gene expression in EpiSCs treated with BRD0705/IWR1, ActivinA, bFGF, and IWR, with normalization to GAPDH. Error bars indicate the SEM from technical triplicates. n=3. (C) qRT-PCR analysis of marker gene expression in EpiSCs maintained in BRD0705/IWR1 and ActivinA/bFGF/XAV939, normalized to GAPDH. Error bars indicate the SEM from technical triplicates, with N=3. (D) Schematic representation of the experimental workflow showing the injection of P10 BRD0705/IWR1-expanded ESCs or EpiSCs into E3.5 blastocysts, followed by embryo development to E10.5 (Left). Representative phase-contrast and fluorescence images of E10.5 mouse embryos derived from blastocyst injection of GFP-expressing ESCs or EpiSCs cultured with BRD0705/IWR1 (right). Scale bars, 500 μm. (E) Shown are phase-contrast and fluorescence images of gonads originating from E15.5 mouse embryos obtained from blastocyst injection of ESCs-GFP cultured with BRD0705/IWR1. Scale bars, 350 μm. (F) Phase-contrast and fluorescence images show #1 embryonic gonad cells (EGCs) derived from the gonads of E15.5 chimeric embryos formed by ESCs-GFP cultured in BRD0705/IWR1. EGCs cultured under 2iL conditions. Scale bars, 250 μm. (G) IF analysis verifies the expression of GFP and OCT4 in EGCs cultured in 2iL. Scale bar 50μm. (H) The schematic illustrates the co-culture of RFP-Zeocin-resistant ESCs and GFP-Puromycin-resistant EpiSCs under BRD0705/IWR1 conditions for 12 passages, followed by selection with Zeocin or Puromycin to isolate ESCs or EpiSCs, respectively. (I) Bright-field and fluorescent images show RFP/GFP expression in co-cultured ESCs (RFP-Zeocin) and EpiSCs (GFP-Puromycin) under BRD0705/IWR1, AFX, and 2iL conditions, following the transition of passage 12 ESCs/EpiSCs maintained in BRD0705/IWR1. Scale bar: 100 μm. (J) The bar graph represents the ratio of ESCs (RFP-Zeocin) and EpiSCs (GFP-Puromycin) shown in Figure I, under BRD0705/IWR1, AFX, and 2iL conditions. (K) Bright-field and fluorescent images show RFP and GFP expression in ESCs (RFP-Zeocin)/EpiSCs (GFP-Puromycin) co-cultured cells after zeocin selection, followed by 3 passages in BRD0705/IWR1, AFX, and 2iL culture conditions. Scale bar: 100 μm. The bar graph represents the proportion of GFP+ or RFP+ cells under different conditions in the panel above. Scale bar: 100 μm. (L) Bright-field and fluorescent images illustrate the RFP and GFP expression in ESCs (RFP-Zeocin)/EpiSCs (GFP-Puromycin) co-cultured cells after puromycin selection, subsequently cultured for 2 passages in BRD0705/IWR1, AFX, and 2iL conditions. Scale bar: 500 μm. The bar graph represents the proportion of GFP+ or RFP+ positive cells under different conditions in the panel above. Scale bar: 100 μm. (M) The schematic shows the generation of chimeric embryos by injecting ESCs (RFP-Zeocin)/EpiSCs (GFP-Puromycin) mixed cells, co-cultured for 14 passages with BRD0705/IWR1, into WT E3.5 embryos, resulting in their contribution to E10.5 embryonic development. (N) Representative morphology of P14 co-cultured ESCs (RFP+) and EpiSCs (GFP+) under BRD0705/IWR1 conditions used for injection into WT E3.5 blastocysts, along with the respective proportions of these two cell types. Scale bar: 100 μm. (O) Images show E10.5 embryos derived from E3.5 blastocyst injection of ESCs (RFP-Zeocin)/EpiSCs (GFP-Puromycin) mixed cells co-cultured in BRD0705/IWR1 for 14 passages, illustrating the distribution and integration of the injected cells in the embryos. Scale bar: 500 μm. (P) Summary of chimera experiments at E10.5.

### ESCs and EpiSCs Maintained in BRD0705/IWR1 Retain Their Respective Pluripotent States

After observing that BRD0705/IWR1 supports the long-term expansion of both ESCs and EpiSCs, we evaluated whether this combination could also preserve the distinct pluripotent states of each cell type. The self-renewal of naive ESCs is typically maintained in 2iL,^9, 43^ whereas primed EpiSCs require AFX conditions.^16^ When cultured in AFX, ESCs convert to EpiSCs and adopt a primed state.^18^ After 25 passages in BRD0705/IWR1, ESCs readily transitioned to 2iL conditions and continued self-renewing. In contrast, transferring EpiSCs from BRD0705/IWR1 to 2iL resulted in cell death or differentiation (Figure 2A). However, both BRD0705/IWR1-expanded ESCs and EpiSCs could be successfully propagated upon transfer to AFX (Figure 2A). These results indicate that BRD0705/IWR1 maintain each cell type in its respective pluripotent state.

Both ESCs and EpiSCs cultured in BRD0705/IWR1 expressed the pluripotency markers NANOG and OCT4 (Figures 2B and S3A) and lacked expression of the primitive streak marker *Foxa2*^44^ and the primitive endoderm marker *Gata4*^45^ (Figure 2B), confirming their pluripotent status. Notably, BRD0705/IWR1-expanded ESCs, similar to 2iL-ESCs, exhibited high levels of the naive pluripotency marker *Rex1*^43^ while showing little to no expression of the primed pluripotency markers *Fgf5*^12^ and *Otx2*^46^ (Figure 2B). After 25 passages in BRD0705/IWR1, ESCs readily transitioned to 2iL and continued to self-renew (Figure 2A). These transitioned cells exhibited gene expression profiles comparable to conventional 2iL-ESCs, further supporting that BRD0705/IWR1 maintains ESCs in a naive pluripotent state (Figure 2B). In contrast, EpiSCs maintained in either BRD0705/IWR1 or AFX displayed high expression of *Fgf5* and *Otx2*, and minimal *Rex1* expression, consistent with a primed identity (Figure 2C). Finally, both BRD0705/IWR1-expanded ESCs and EpiSCs retained the ability to differentiate into all three germ layers: ectoderm, mesoderm, and endoderm, *in vitro* (Figure S3B). Collectively, these findings demonstrate that BRD0705/IWR1 effectively maintains the distinct pluripotent states of ESCs and EpiSCs.

Chimera formation remains the gold standard for demonstrating naive pluripotency. When injected into mouse blastocysts, BRD0705/IWR1-expanded ESCs efficiently contributed to chimeric embryos, whereas BRD0705/IWR1-expanded EpiSCs did not (Figures 2D, S3C-F). To further assess the developmental potential of BRD0705/IWR1-expanded ESCs, we evaluated their ability to contribute to the germ cell lineage. Gonads were isolated from E15.5 chimeric embryos generated using GFP-labeled BRD0705/IWR1-expanded ESCs (Figure 2E). Of four gonads collected from three chimeric embryos, two—each from a different embryo— contained GFP-positive cells. To determine whether these GFP+ cells were primordial germ cells (PGCs), the gonads were dissociated and cultured under 2iL conditions^47, 48^. GFP+ embryonic germ (EG) cell colonies emerged from both gonads and expanded robustly (Figures 2F and S3E). These EG colonies co-expressed OCT4 and GFP (Figure 2G), indicating that BRD0705/IWR1-expanded ESCs contributed to the germline in chimeric embryos.

In summary, ESCs maintained long-term in BRD0705/IWR1 retained naive pluripotency, as demonstrated by marker expression and functional contributions to both chimera and germ cell lineages. In contrast, EpiSCs expanded under BRD0705/IWR1 retained a primed pluripotent state, supported by both molecular profiling and lack of chimeric contribution.

### ESC/EpiSC Co-cultures Maintain Cell Identities Following Long-Term Exposure to BRD0705/IWR1 Conditions

Notably, the same culture condition, BRD0705/IWR1, supported the maintenance of two distinct pluripotent states. To rigorously assess whether ESCs and EpiSCs retained their respective identities during extended co-culture in BRD0705/IWR1, RFP-labeled ESCs and GFP-labeled EpiSCs were mixed at a 1:1 ratio and cultured together for 12 passages (over one month) (Figure 2H). To evaluate whether these long-term co-cultured cells preserved their original characteristics, passage 12 (P12) co-cultured cells were replated separately under BRD0705/IWR1, AFX, and 2iL conditions (Figures 2I and 2J). Under continued BRD0705/IWR1 conditions, ESCs and EpiSCs remained an approximate 1:1 ratio (Figures 2I and 2J). By contrast, in AFX conditions, the proportion of EpiSCs-derived cells increased significantly to 67.5%, while ESC-derived cells accounted for 32.5%. Conversely, in 2iL conditions, 95.5% of the population consisted of ESC-derived cells (Figures 2I and 2J). These results strongly support that BRD0705/IWR1 simultaneously supports the self-renewal of both ESCs and EpiSCs without inducing interconversion or dominance of one state over the other.

To further assess whether ESCs and EpiSCs retained their intrinsic properties after long-term co-culture, passage 13 cells were subjected to selection using zeocin and puromycin. Zeocin-selected RFP+ ESCs exhibited robust growth under BRD0705/IWR1, AFX, and 2iL conditions (Figure 2K). In contrast, puromycin-selected GFP+ EpiSCs grew stably under BRD0705/IWR1 and AFX but failed to survive in 2iL conditions (Figure 2L), further supporting their retention of a primed identity.

After 14 passages (over one month), the mixed population (43.4% RFP+ ESCs, 56.6% GFP+ EpiSCs) cultured under BRD0705/IWR1 was injected into mouse blastocysts to assess lineage contribution in vivo (Figures 2M and 2N). Notably, only RFP+ ESCs contributed to E10.5 chimeras (55.6%), while GFP+ EpiSC-derived cells were not detected (Figures 2O and 2P), consistent with their limited developmental potential.

Taken together, these findings demonstrate that ESCs and EpiSCs co-cultured under BRD0705/IWR1 conditions maintain their distinct molecular identities and developmental potentials, validating this culture system as a robust platform for sustaining heterogeneous pluripotent states in parallel.

### BRD0705/IWR1 Uniquely Preserves the Transcriptomic and Epigenetic Identities of ESCs and EpiSCs

To assess whether BRD0705/IWR1 preserves the identities of ESCs and EpiSCs after extended co-culture, we performed single-cell RNA sequencing (scRNA-seq) on RFP⁺ ESCs and GFP⁺ EpiSCs following 16 passages under BRD0705/IWR1 conditions (Figure 3A). Uniform manifold approximation and projection (UMAP) analysis revealed three distinct clusters corresponding to RFP+ ESCs, GFP+ EpiSCs, and GFP-/RFP-/Vimentin+ feeder cells (Figure 3B). Examination of naive and primed pluripotency marker genes within these clusters showed that the core pluripotency marker *Nanog* was expressed in both GFP+ and RFP+ cells, indicating that pluripotency was maintained across both populations (Figure 3C). Naive pluripotency markers, including *Esrrb, Zfp42, Nr0b1, Tfcp2l1, Tbx3, Tcl1, and Prdm14*, were predominantly expressed in the RFP+ ESC cluster, while the markers for primed state such as *Fgf5* and *Pitx2* were largely restricted to the GFP+ EpiSC cluster (Figure 3C).

**Figure 3.**
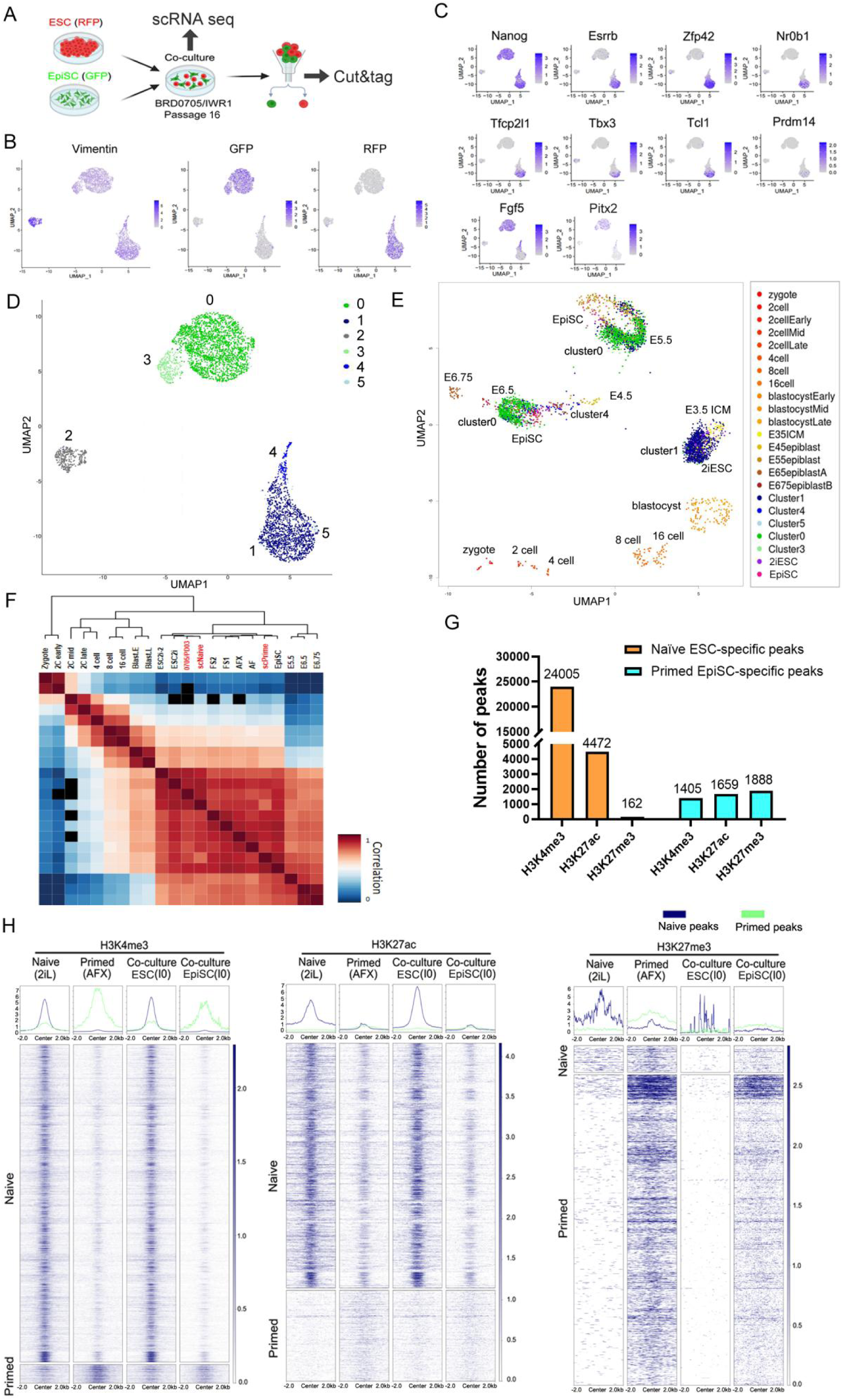
scRNA seq and Chromatin Landscape Analysis. (A) Schematic illustration of the experimental design. Naive ESCs (RFP+) and EpiSCs (GFP+) were co-cultured under BRD0705/IWR1 conditions for 16 passages, followed by scRNA-seq and Cut&tag analyses. (B) UMAP plots displaying the expression of Vimentin, GFP, and RFP confirm the presence of ESCs and EpiSCs in co-culture under BRD0705/IWR1 conditions. (C) UMAP visualization of key pluripotency-related genes, including the PSC marker Nanog; ESC markers (*Esrrb, Zfp42, Nr0b1, Tfcp2l1, Tbx3, Tcl1, and Prdm14*); and the EpiSCs marker *Fgf5* and *Pitx2*, highlights the transcriptional differences between ESCs and EpiSCs. (D) UMAP clustering of single-cell transcriptomes reveals distinct cell populations within the co-culture cells, labeled as clusters 0–5. (E) Clusters 1–5 from (D) were projected onto a UMAP plot of early embryonic developmental stages. (F) Heatmap showing the transcriptomic correlation between co-cultured cells under BRD0705/IWR1 conditions and reference ESCs^50^, EpiSCs^50^, and formative cells^16^ cultured under different conditions, indicating their resemblance to naive and primed pluripotent states. scNaive: single-cell sequencing results of naive ESCs in BI co-cultured cells. scPrime: single-cell sequencing results of primed PSCs in BI co-cultured cells. (G) Bar graph quantifying the number of naive ESCs-specific and primed EpiSCs-specific chromatin accessibility peaks for H3K4me3, H3K27ac, and H3K27me3 modifications. (H) Heatmaps displaying Cut&Tag profiles for H3K4me3, H3K27ac, and H3K27me3 modifications in naive ESCs (2iL), primed EpiSCs (AFX), and BRD0705/IWR1 co-cultured ESCs (RFP+) and EpiSCs (GFP+). Naive ESCs exhibit distinct chromatin accessibility patterns compared to primed EpiSCs, and co-cultured ESCs and EpiSCs maintain their respective epigenetic signatures, confirming the long-term maintenance of their identities under BRD0705/IWR1 conditions.

GFP+ and RFP+ cells were further classified into subpopulations based on gene expression clustering, with cluster 0 and cluster 1 representing the major groups of GFP+ and RFP+ cells, respectively (Figure 3D). UMAP analysis of scRNA-seq data from co-cultured cells treated with BRD0705/IWR1 was performed and compared to reference profiles from embryos spanning from the zygote to E6.75 embryo stages. RFP+ ESCs cultured long-term under BRD0705/IWR1 conditions predominantly mapped to the E3.5 stage, consistent with the distribution of naive ESCs cultured in 2iL. In contrast, GFP+ EpiSCs primarily mapped to E5.5 and E6.5, mirroring the distribution of EpiSCs cultured in ActivinA/bFGF (AF) (Figure 3E). In addition, unsupervised hierarchical clustering analysis showed that ESCs and EpiSCs co-cultured under BRD0705/IWR1 exhibited transcriptomic profiles closely correlated with ESCs and EpiSCs cultured in 2i and Activin A/bFGF (AF) (Figures 3F and S3I). These findings suggest that even after extended *in vitro* co-culture under BRD0705/IWR1 conditions, ESCs and EpiSCs preserved their distinct developmental identities and lineage-appropriate transcriptional programs.

In addition to gene expression analysis, the epigenetic states of co-cultured cells were evaluated. State-specific peaks for histone marks H3K4me3, H3K27ac, and H3K27me3 were first defined in naive ESCs and primed EpiSCs using publicly available data^16^ (Figure 3G). Cut&Tag^49^ was then performed on cells co-cultured under BRD0705/IWR1 conditions to assess the occupancy of H3K4me3, H3K27ac, and H3K27me3. Distinct and highly specific peak profiles for each histone mark were observed between ESCs and EpiSCs. Moreover, the unique peaks exhibited by each population closely resembled those of naive ESCs and primed ESCs, respectively, indicating that chromatin state at promoter and enhancer regions remained cell-type specific and similar to that of traditionally cultured naive and primed ESCs (Figure 3H). These findings demonstrated that BRD0705/IWR1 stably preserved the epigenetic landscapes of ESCs and EpiSCs during long term culture, providing a robust molecular foundation for the maintenance of distinct pluripotent identities.

### BRD0705 selectively inhibits GSK3α to promote self-renewal of ESCs and EpiSCs

While IWR1 is a well-characterized tankyrase inhibitor known to play a role in the self-renewal of pluripotent stem cells^12^, the role of BRD0705 in stem cell maintenance remains largely unexplored. To determine whether BRD0705 promotes the self-renewal of ESCs and EpiSCs through selective inhibition of GSK3α, we examined its effect in *Gsk3α/β* double knockout (DKO) ESCs^22^. In serum-free N2B27 medium, the self-renewal of DKO ESCs was sustained by PD03 alone, and the addition of either BRD0705 or the pan-GSK3 inhibitor CHIR provided no further benefit (Figures 4A, 4B, S4A and S4B).

**Figure 4.**
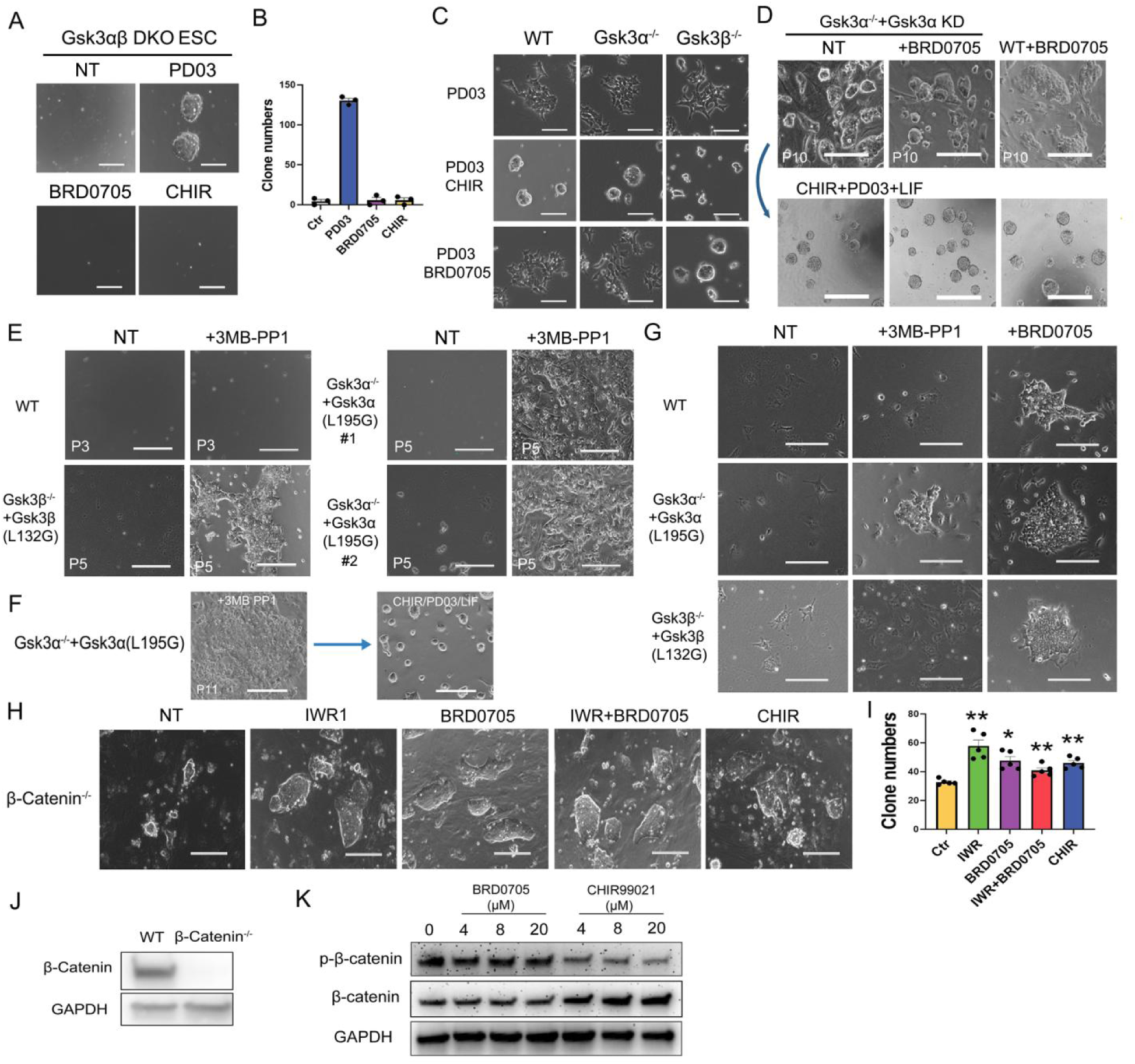
BRD0705 enhances stemness maintenance in other types of stem cells. (A) Morphological comparison of #1 Gsk3αβ DKO ESC line under different treatment conditions. Cells were cultured in non-treated control conditions, PD03, BRD0705, and CHIR. Cell cultured in DMEM/FBS medium for Passage 3. Scale bars represent 100 μm. (B) Quantification of clone numbers corresponding to the images in Figure A. PD03 treatment significantly increases *Gsk3α/β* DKO ESCs clone numbers compared to control and other conditions (BRD0705 and CHIR), which show minimal clone formation. Error bars indicate mean ± SEM. (C) Morphology of WT, *Gsk3α^-/-^*, and *Gsk3β^-/-^* ESCs treated with PD03, PD03+CHIR, or PD03+BRD0705. PD03+CHIR induces ‘dome-like’ colonies in WT, *Gsk3α^-/-^* and *Gsk3β^-/-^*. PD03+BRD0705 induces ‘dome-like’ colony formation only in *Gsk3β^-/-^* cells, while other genotypes show a spread-out morphology. Cell cultured in N2B27 for day 4. Scale bars, 200 µm. (D) The top row shows *Gsk3α^-/-^-Gsk3αKD* ESCs under NT, BRD0705 condition, and WT ESCs treated with BRD0705 after 10 passages (P10). The bottom row displays the morphology of the corresponding P10 cells after being transitioned to 2iL conditions for 3 passages. Scale bars represent 100 µm. (E) Survival of ESCs lines under different conditions. Representative images of WT, *Gsk3α^−/−^+Gsk3α-L195G*, and *Gsk3β^−/−^+Gsk3β-L132G* ESCs cultured with or without 3MB-PP1. #1 and #2 represent two different cell lines. WT ESCs displayed complete cell death by passage 3 (P3) under both conditions, while Gsk3-modified ESCs maintained viability at passage 5 (P5) with visible cell colonies. Scale bars represent 200 μm. (F) Morphology of *Gsk3α^−/−^+Gsk3α-L195G* ESCs under different culture conditions. Representative images of cells at passage 11 (P11) cultured in +3MB-PP1 (left) and following a switch to 2iL (right). Scale bars, 200 μm. (G) Morphology of WT, *Gsk3α^−/−^+Gsk3α-L195G*, and *Gsk3β^−/−^+Gsk3β-L132G* EpiSCs under different treatments. Representative images show cells cultured in DMEM/FBS medium for 3 passages in NT, +3MB-PP1, and +BRD0705 conditions. Scale bars, 200 μm. (H) Morphological analysis of β-Catenin^-/-^ ESCs cultured in LIF/N2B27 on feeder under different treatment conditions. Representative images display cells cultured in NT, IWR1, BRD0705, IWR+BRD0705, and CHIR conditions. Scale bars, 100 μm. (I) Quantification of Figure I. Clone numbers of β-Catenin^-/-^ ESCs under different treatments: Ctr, IWR1, BRD0705, IWR1/BRD0705, and CHIR. Data are presented as mean ± SEM, *, p < 0.05, **, p < 0.01. (J) Western blot analysis of β-Catenin in WT and *β-Catenin^-/-^* ESCs. β-Catenin is detected in WT but absent in β-Catenin^-/-^ cells. GAPDH serves as the control. (K) Western blot analysis of phosphorylated β-catenin (p-β-catenin), total β-catenin, and GAPDH in response to different concentrations (0, 4, 8, 20 µM) of BRD0705 and CHIR.

Next, *Gsk3α*^⁻/⁻^ and *Gsk3β*^⁻/⁻^ ESCs were used to determine whether BRD0705 selectively inhibits GSK3α. When combined with PD03 in N2B27, pan-GSK3 inhibition using CHIR maintained self-renewal in wild-type, *Gsk3α*^⁻/⁻^, and *Gsk3β*^⁻/⁻^ ESCs (Figures 4C, S4D). Notably, under serum-free PD03/N2B27 conditions, the addition of BRD0705 was sufficient to maintain self-renewal in *Gsk3β*^⁻/⁻^ ESCs but not in *Gsk3α*^⁻/⁻^ ESCs (Figures 4C, S4C, and S4D), demonstrating that BRD0705 promotes ESC self-renewal through selective inhibition of GSK3α, rather than GSK3β.

To further validate that BRD0705 acts via inhibition of GSK3α, we used ESCs in which endogenous GSK3α was replaced with a kinase-dead (KD) mutant (*GSK3α-K148R*).^22, 51^ This approach allowed specific disruption of kinase activity without affecting other GSK3α functions such as heterodimerization, which are lost in *Gsk3α^-/-^* ESCs. Under DMEM/10% FBS conditions without additional inhibitors, *Gsk3α^-/-^*+*Gsk3α-KD* ESCs remained undifferentiated, whereas WT ESCs either differentiated or died by the third passage (Figure S4E). Next, both WT ESCs cultured with BRD0705 and Gsk3α^-/-^+Gsk3α-KD ESCs cultured without BRD0705 were passaged for ten generations and then transferred to 2iL conditions. In both cases, the cells remained undifferentiated in 2iL (Figure 4D), strongly indicating that selective inhibition of GSK3α kinase activity is sufficient to maintain ESC self-renewal.

To independently confirm this conclusion, we employed a chemical-genetic strategy using analog-sensitive GSK3 mutants: *GSK3α-L195G* and *GSK3β-L132G*. These mutants specifically bind the “bumped” kinase inhibitor 3MB-PP1, thereby recapitulating the kinase-inhibition observed with small-molecule-specific inhibition of GSK3α/β.^22^ Selective inhibition of GSK3α in *Gsk3α^−/−^+ Gsk3α-L195G* ESCs^22^ by 3MB-PP1 was sufficient to support self-renewal (Figure 4E), further confirming the role of GSK3α kinase inhibition in promoting ESC self-renewal. Interestingly, selective inhibition of GSK3β also supported ESC self-renewal (Figure 4E). *Gsk3α^−/−^+Gsk3α-L195G* ESCs maintained for 11 passages with 3MB-PP1 were readily reverted to 2iL conditions (Figure 4F), indicating the retention of their naive pluripotent state. Inhibition of GSK3α, whether by 3MB-PP1 or BRD0705, also promoted EpiSC self-renewal, whereas selective inhibition of GSK3β led to differentiation of EpiSCs (Figure 4G). CHIR maintains ESCs self-renewal by stabilizing β-catenin through the inhibition of GSK3.^9^ Therefore, we aimed to investigate whether GSK3α exerts its effects through a similar β-catenin-mediated mechanism. Unexpectedly, BRD0705 maintains its ability to promote self-renewal in ESCs even in the absence of β-catenin (Figures 4H-4J). Unlike CHIR, BRD0705 does not regulate the phosphorylation of β-catenin in ESCs (Figure 4K). Collectively, these findings demonstrate that BRD0705 facilitates ESC self-renewal primarily through selective inhibition of GSK3α kinase activity, independent of β-catenin signaling. This identifies GSK3α as a novel molecular target for the maintenance of pluripotency and suggests the existence of a yet undescribed downstream signaling mechanism distinct from canonical WNT/β-catenin pathways.

### BRD0705 Promotes Self-Renewal of Formative Pluripotent Stem Cells and Neural Stem Cells

Naive ESCs correspond to the cellular stage of the inner cell mass of a mouse embryo at E3.5–E4.5.^43^ In contrast, EpiSCs exhibit phenotypes characteristic of the mid-gastrula stage, closely resembling the primed epiblast of the anterior primitive streak.^11, 52^ Formative stem cells represent an intermediate state, corresponding to the post-implantation epiblast at the E5.5– E6.0 stage, characterized by transient presence, heterogeneity, and enrichment in cells related to the pre-streak epiblast.^14, 53^

Given BRD0705’s role in maintaining the pluripotency of both ESC and EpiSC, we investigated its potential to support the self-renewal of formative stem cells. Typically maintained under Activin A/XAV939/BMS493 (A_lo_XR)^16^ conditions, mouse formative stem cells were successfully transitioned to BRD0705/IWR1 conditions while retaining their formative pluripotent state (Figure S5A). After long-term culture in BRD0705/IWR1, these cells could be transferred back to A_lo_XR conditions, where they continued to maintain their formative state (Figure S5A). Formative cells cultured under BRD0705/IWR1 conditions exhibited a gene expression profile comparable to those maintained in A_lo_XR conditions (Figure S5B).

To test its effect beyond pluripotent stem cells, the ability of BRD0705 to sustain neural stem cell stemness was assessed. SOX1+ neural stem cells were isolated and expanded under bFGF and EGF^54, 55^, either with or without BRD0705 (Figure S5C). Notably, cells treated with BRD0705 retained SOX1 expression for an extended period and across multiple passages compared to the control group (Figures S5C, S5D, and S5E). Collectively, these findings establish that BRD0705 promotes the long-term maintenance of not only ESCs and EpiSCs, but also formative stem cells and neural stem cells, through selective inhibition of GSK3α. This identifies GSK3α as a novel target for stem cell maintenance and implicates a fundamental, previously undescribed signaling mechanism that supports self-renewal across diverse stem cell types.

## Discussion

In this study, we identified GSK3α as a key regulator of self-renewal and pluripotency in mouse pluripotent stem cells across multiple states. Our findings demonstrate that selective inhibition of GSK3α with BRD0705 stabilizes pluripotent stem cells within their current identity, preventing both transition between pluripotent states and exit from pluripotency. This effect is evident in both naive ESCs and primed EpiSCs, which remain stably naive or primed, respectively, under BRD0705/IWR1 culture conditions. We propose that GSK3α acts as a stemness checkpoint in mouse pluripotent stem cells, with its inhibition insulating stem cells from differentiation cues and thereby enhancing their intrinsic self-renewal capacity, a role that has been previously underexplored in the regulation of stem cell fate.

One of the most striking findings is that BRD0705 supports the self-renewal of both naive ESCs and primed EpiSCs. When combined with IWR1, BRD0705 enables the long-term expansion of both cell types while preserving their distinct molecular identities. ESCs cultured in BRD0705/IWR1 readily transition back to 2iL conditions while maintaining naive markers or functionality, whereas EpiSCs maintained in BRD0705/IWR1 can readily revert to AFX conditions but not to 2iL, reflecting their preserved primed pluripotency identity. These results highlight the specificity of BRD0705 in maintaining pluripotent states and suggest that GSK3α functions as a molecular gatekeeper, whose inhibition prevents differentiation through a mechanism independent of the β-catenin pathway.

Specific inhibition of GSK3α by BRD0705 supports ESC self-renewal in the presence of either serum or feeder cells. However, BRD0705 alone is not sufficient to maintain the undifferentiated state; ESCs cultured under serum-free, feeder-free conditions undergo neural differentiation despite selective inhibition of GSK3α.^22^ This context-dependent effect mirrors that of LIF, which supports ESC self-renewal in serum-containing environments but fails to do so under defined, serum-free, feeder-free conditions, where ESCs also preferentially differentiate toward the neural lineage, even in the continued presence of LIF.^56, 57^ These parallels support a model in which the ability of GSK3α inhibition, like LIF-mediated STAT3 activation, to sustain self-renewal is contingent upon extrinsic cues such as serum components or feeder-derived factors. Thus, the extracellular environment plays a critical role in modulating the cellular response to kinase inhibition and directing stem cell fate. Our findings underscore the importance of contextual signals in determining whether GSK3α inhibition promotes self-renewal or lineage commitment.

Although GSK3α is highly conserved among mammals, its absence in avian species^58^ suggests that it may serve a mammal-specific role in regulating pluripotency. Our findings reveal that, unlike GSK3β, GSK3α promotes PSCs self-renewal through a β-catenin– independent mechanism. This distinction highlights that BRD0705 and CHIR operate through fundamentally different pathways to maintain ESC self-renewal. This distinction has significant implications for understanding species-specific mechanisms of stem cell regulation. In rodent ESCs, nuclear translocation of β-catenin promotes self-renewal, while in human naive ESCs, it drives differentiation.^59^ Reflecting this divergence, recently developed culture conditions for maintaining naive human ESCs (PXGL)^60^ and chimera-competent monkey ESCs (4CL)^61^ intentionally exclude CHIR and instead incorporate WNT pathway inhibitors such as IWR1. These observations underscore the limitations of β-catenin–dependent pathways in supporting naive pluripotency across species. Therefore, the identification of BRD0705 as a β-catenin– independent GSK3α inhibitor offers a significant advance. It provides a promising tool for developing ESC culture systems applicable to a broad range of mammalian species, potentially enabling the derivation and maintenance of naive ESCs in contexts where canonical WNT signaling is incompatible with self-renewal.

Our findings also extend beyond pluripotent stem cells. BRD0705 not only promotes the self-renewal of formative pluripotent stem cells, an intermediate state between naive and primed pluripotency^16^, but also supports the maintenance of neural stem cell. BRD0705 significantly increased neural stem cell numbers and enabled long-term maintenance of SOX1^62^ expression (Figure S5 C-E). This suggests that GSK3α’s role as a stemness checkpoint may apply broadly to other tissue-specific stem cells. The implications of this are significant, as GSK3α inhibition could provide a novel approach to enhance the self-renewal and maintenance of various stem cell types, potentially facilitating regenerative therapies and stem cell-based models of development and disease.

Our findings raise intriguing questions about the existence of other stemness checkpoint proteins that might similarly prevent cell state transitions and maintain stemness across diverse cell types. Identifying additional proteins with functions analogous to GSK3α could broaden our understanding of the regulatory mechanisms that preserve stem cell identity and pluripotency. This approach could lead to the discovery of universal conditions for maintaining stemness, facilitating stem cell research and applications across species.

In conclusion, our study positions GSK3α as a central regulator of stem cell pluripotency and suggests that its inhibition offers a promising strategy for sustaining stemness in both pluripotent and tissue-specific stem cells. Future studies may identify more such checkpoint proteins, paving the way for robust, species-independent culture conditions that maintain stem cell states for various biomedical applications.

### Limitations of the study

Our study demonstrates that selective inhibition of GSK3α plays a critical role in maintaining the stemness of both ESC and EpiSC. However, the precise molecular mechanisms underlying this effect require further investigation. We showed that BRD0705 preserves ESC stemness in a β-catenin–independent manner, suggesting that GSK3α may regulate stem cell maintenance through a novel signaling pathway. Further elucidating this mechanism will enhance our understanding of pluripotency regulation. Additionally, it remains to be determined whether GSK3α inhibition consistently sustains stemness across different pluripotent states and whether the ability of BRD0705/IWR1 to maintain both ESC and EpiSC identities extends to other species.

## METHODS

### Key resources table

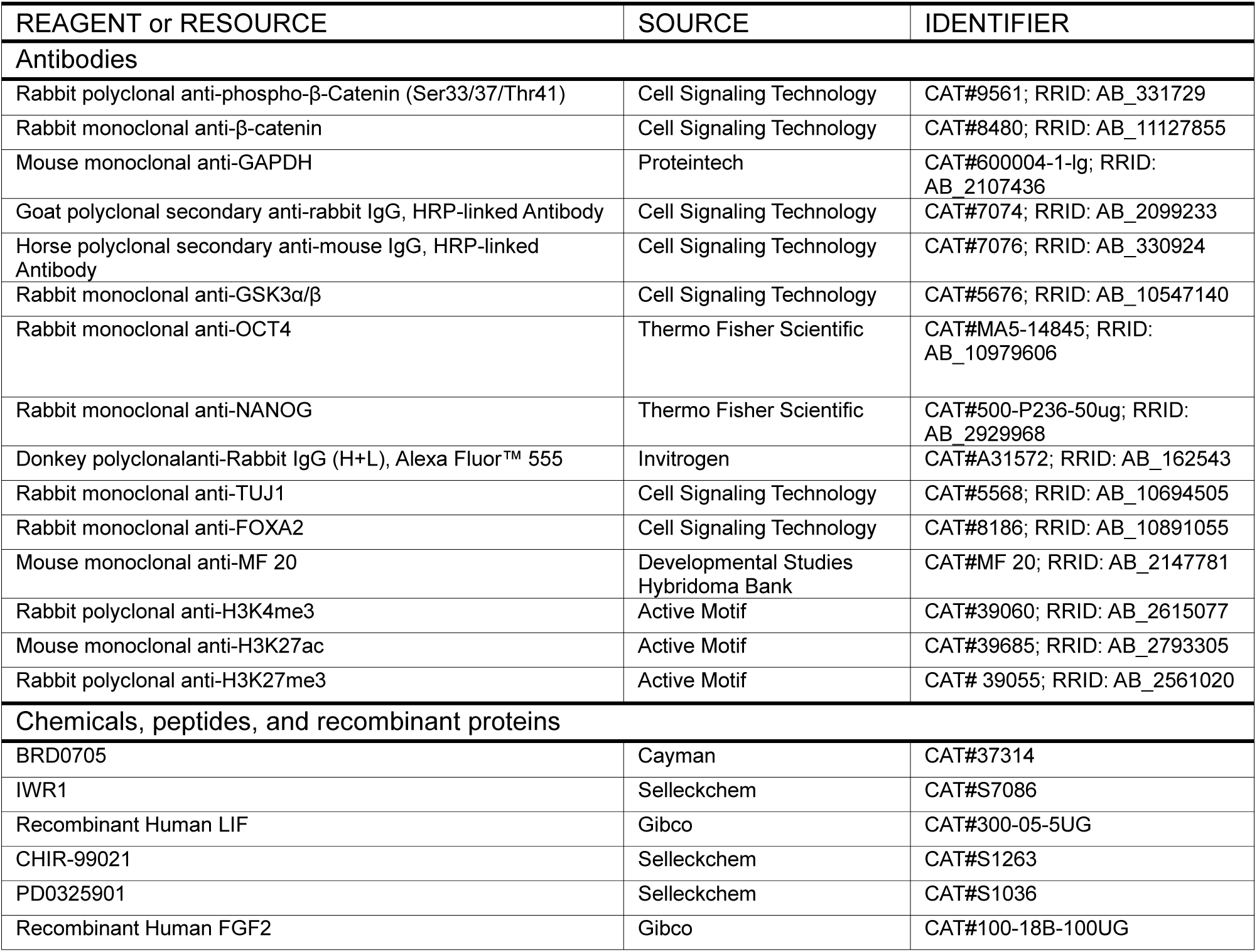
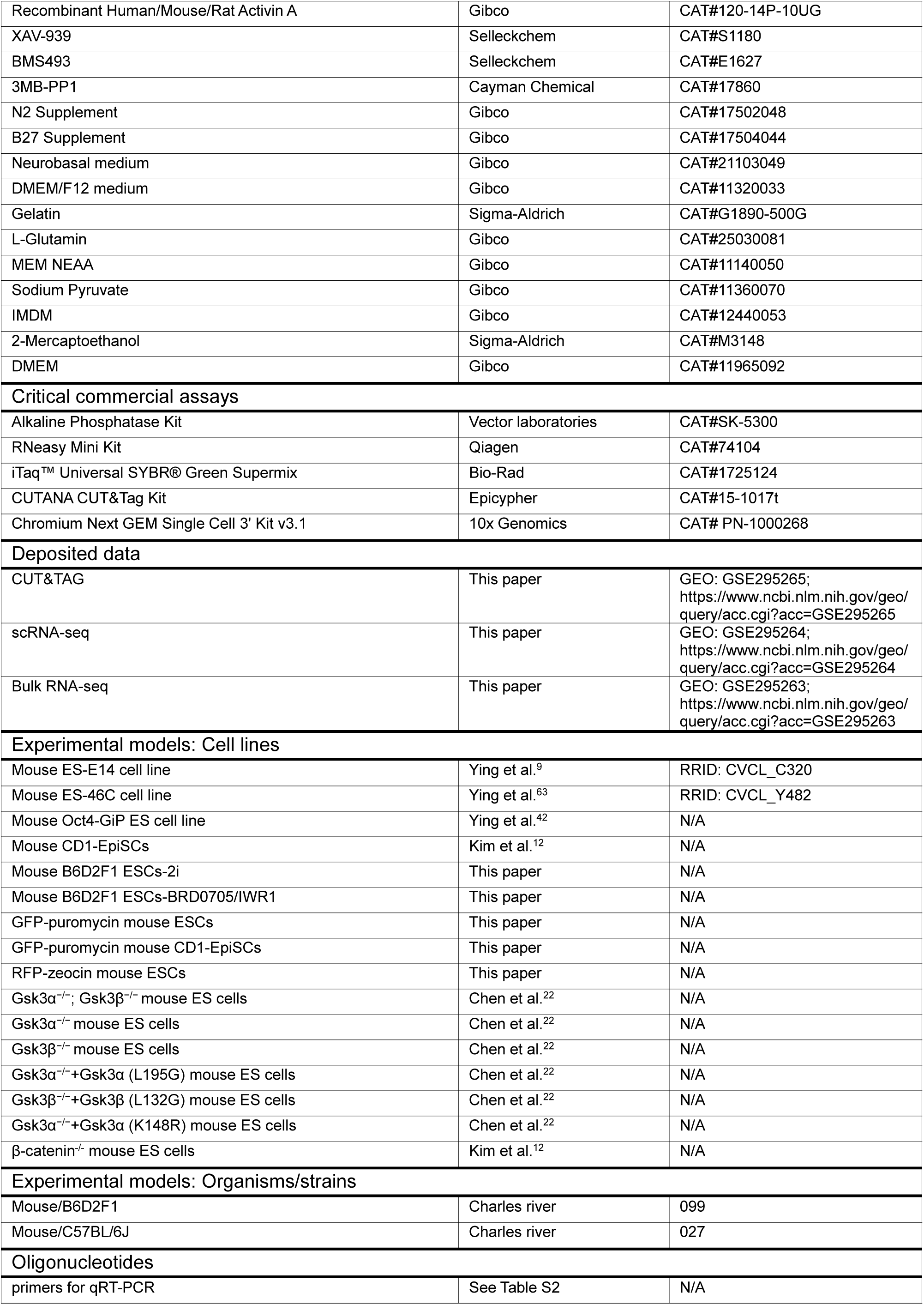
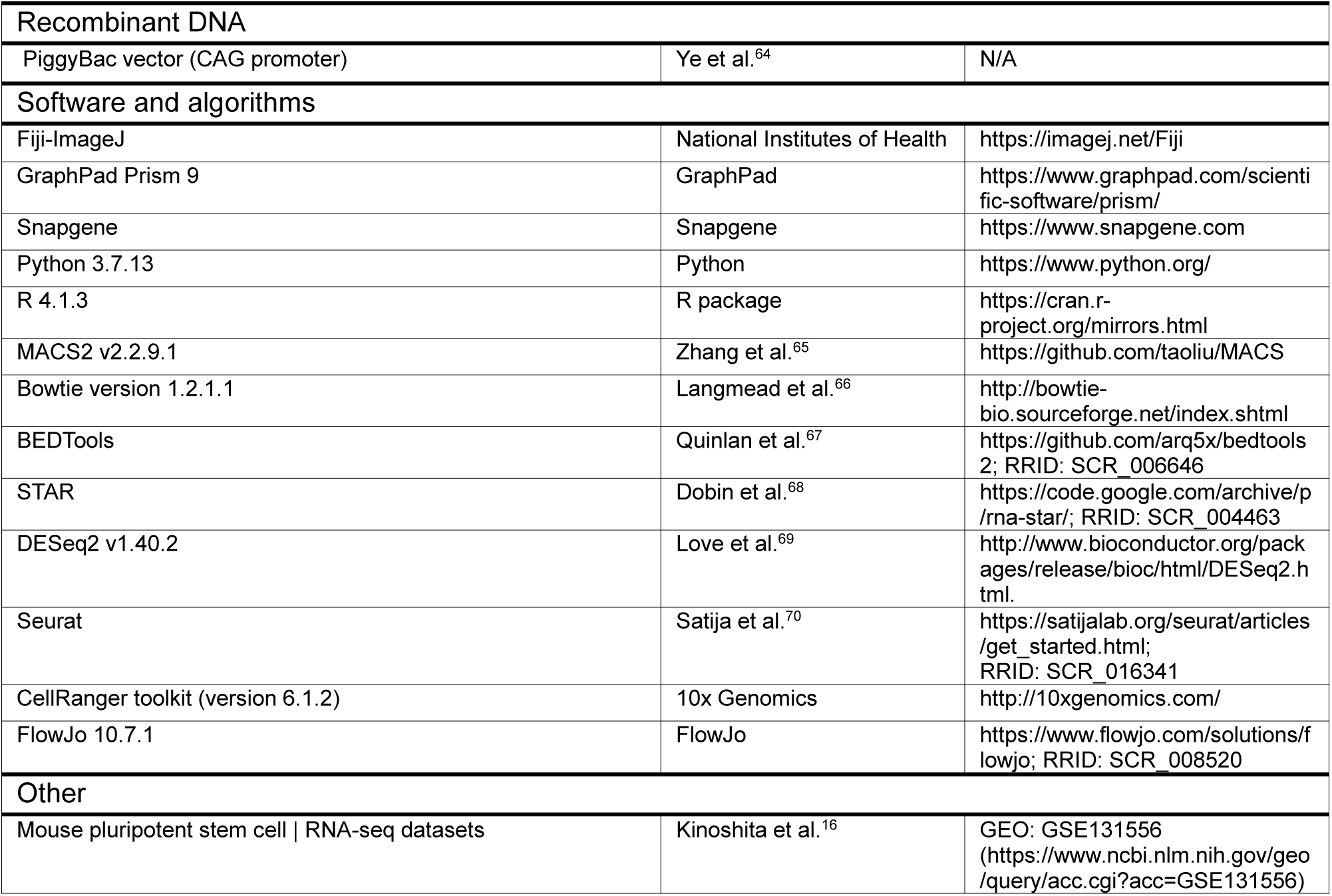

## RESOURCE AVAILABILITY

### Lead Contact

Further information and requests for resources and reagents should be directed to and will be fulfilled by the Lead Contact, Qilong Ying

### Materials Availability

All reagents and stable cell lines related to this manuscript are available upon request from the Lead Contact, contingent on the completion of a Materials Transfer Agreement with the University of Southern California.

### Data and Code Availability

Raw sequencing data and code will be deposited upon publication.

## EXPERIMENTAL MODEL AND SUBJECT DETAILS

### Mice

Adult female mice were utilized in these experiments. Embryos for cell line derivation were obtained from B6D2F1 strains. C57BL/6J mice provided host embryos for chimera generation, and ARC mice were used as recipients for embryo transfer. The experiments involving the injection of embryonic stem cells into blastocysts, blastocyst transplantation, and the culture of mice after embryo transfer were all conducted at the Irvine Transgenic Mouse Facility at the

University of California, Irvine. All studies following guidelines for laboratory animal care and use, and were carried out under the authority of IACUC protocol #AUP-22-126. The experiments related to obtaining mouse embryos for ESC extraction were conducted at the Department of Animal Resources facility at the University of Southern California. The project received approval from the Animal Welfare and Ethical Review Body at the University of California, Irvine and University of Southern California.

### Cell Cultures

Cell lines are detailed in the Key Resources Table. They were cultured in the incubators at 37°C with 5% CO₂, without the use of antibiotics.

### Conversion of ESCs to Formative cells

ESCs were initially plated in A_lo_XR (3 ng/mL activin A, 2 mM XAV939, and 1.0 mM BMS493) in N2B27 medium.^16^ Complete conversion typically requires approximately 4–5 passages. After conversion, the cultures were maintained at higher densities and passaged using accutase.

### Conversion of ESCs to EpiSCs

ESC was initially plated in AFX medium (20 ng/ml Activin A, 20 ng/ml bFGF, and 2 µM XAV-939) in N2B27 medium or DMEM/10%FBS medium. Complete conversion typically requires approximately 4-5 passages.

### Derivation of ESCs from mouse blastocyst

The E3.5 B6D2F1 mouse blastocyst was flushed from the mouse oviduct. The zona pellucida was removed using acid Tyrode’s solution, after which the blastocyst was transferred onto a feeder-coated culture plate. The embryos were cultured in N2B27 medium supplemented with BRD0705/PD03 or CHIR/PD03. After approximately 4-5 days of growth, the blastocysts were digested into single cells using 0.025% trypsin for passaging.

## METHOD DETAILS

### Western blot

The cells were lysed using ice-cold RIPA buffer, comprising 1% Triton X-100, 50 mM Tris, 150 mM NaCl, 0.1% SDS, and 1% sodium deoxycholate, with protease and phosphatase inhibitors (Roche) added. After denaturation at 95°C, the proteins, mixed with Laemmli sample buffer (Bio-Rad), were separated on a 10-15% polyacrylamide gel and transferred onto a PVDF membrane (Millipore) via electrophoresis. The membrane was rinsed with TBST (0.1% Tween-20 in TBS), blocked with 5% Blocker (Bio-Rad) in TBST for 1 hour at room temperature, then incubated with primary antibodies at 4°C overnight, followed by a 1-hour incubation with HRP-conjugated secondary antibodies at room temperature the following day.

### Quantitative real-time PCR (qRT-PCR)

Total RNA was isolated using the RNeasy Mini Kit (Qiagen) following the manufacturer’s instructions. Complementary DNA (cDNA) synthesis was carried out with the iScript cDNA Synthesis Kit (Bio-Rad). Quantitative real-time PCR was conducted using the iTaq Universal SYBR® Green Supermix (Bio-Rad) on a Viia 7 real-time PCR system. Gene expression levels were normalized to Gapdh.

### ESCs culture

The ESCs were cultured on 0.1% gelatin-coated dishes at 37°C in a 5% CO_2_ atmosphere. The DMEM/10% FBS medium was prepared by supplementing DMEM (Gibco) with 10% FBS (Gibco), 0.1 mM β-mercaptoethanol (Sigma), 1% MEM Non-Essential Amino Acids Solution (Gibco), and 2 mM L-glutamine (Gibco). DMEM/F12-N2 medium was prepared by adding 1 ml of N2 (Gibco) 100× stock solution to 100 ml of DMEM/F12. For Neurobasal/B27 medium, 2 ml of B27 (Gibco) and 2 mM L-glutamine were added to 100 ml of Neurobasal medium. N2B27 medium was prepared by mixing DMEM/F12-N2 medium with Neurobasal/B27 medium at a 1:1 ratio, followed by the addition of 0.1 mM β-mercaptoethanol (Sigma). To expand naive-state ESC, 3 µM CHIR-99021 (Selleckchem), 1 µM PD03 (Selleckchem), and 20 ng/ml LIF (Peprotech) (2iL) were added to the DMEM/FBS or N2B27 medium. E14TG2a ESC can be cultured under 2iL conditions in either DMEM/FBS or N2B27 medium, with or without feeders. However, for B6D2F1 ESC, it is necessary to culture them on feeder-coated plates using 2iL/N2B27 or 2iL/DMEM/10% FBS medium. For BRD0705/IWR conditions, 8 µM BRD0705 (Cayman Chemical) and 2.5 µM IWR-1 (Selleckchem) were added to the DMEM/FBS or N2B27 medium. Under BRD0705 only or BRD0705/IWR-1 conditions, ESC require feeder cells when cultured in N2B27 medium, whereas in DMEM with 10% FBS, feeder cells are not required, and pre-coating the culture plates with 0.1% gelatin is sufficient. Ctnnb1^-/-^ ESC^12^ were cultured on γ-irradiated CF-1 MEF feeders in N2B27 medium containing 20 ng/ml LIF and 1 µM PD03 (Selleckchem).

### EpiSCs culture

CD1 EpiSCs were established from the epiblasts of E5.75 CD1 mouse embryos, as previously described.^71^ To generate E14TG2a-EpiSC, E14TG2a-ESC were cultured in a basal medium containing 20 ng/ml Activin A (PeproTech), 20 ng/ml bFGF (PeproTech), and 2 µM XAV-939 (Sigma). Both CD1 and E14TG2a-EpiSC were subsequently maintained in DMEM/10%FBS medium or N2B27 medium with the addition of 1.5 µM CHIR99021 and 2.5 µM IWR-1. When using N2B27 medium, culturing EpiSCs on feeder cells is more effective. However, when using DMEM/FBS medium, feeder cells are not required, and pre-coating the culture plates with 0.1% gelatin is sufficient.

### Chimeras

Approximately 15 individually dissociated cells were injected into each blastocyst-stage embryo. The embryos were then either transferred into pseudo-pregnant mice. Imaging of E10.5 mid-gestation embryos was performed using a Keyence BZ-X800 fluorescence microscope. For the isolation of PGCs from chimeric mice embryos, embryos at E15.5 stage of development are used.

### Embryoid body formation and differentiation

A total of 2×10^4 cells were seeded into AggreWell™400 (STEMCELL Technologies) using either N2B27 medium or IMDM/FBS medium^71^ (IMDM supplemented with 15% fetal bovine serum, 2 mM L-glutamine, 0.05 mg/mL ascorbic acid, and 0.001% monothioglycerol). After 2–3 days of incubation, the resulting embryoid bodies (EBs) were transferred onto laminin- or gelatin-coated plates with fresh medium for outgrowth. For neural induction, ESCs or EpiSCs were plated on laminin-coated plates in N2B27 medium.^63, 72^ For mesoderm induction, ESCs or

EpiSCs were plated on gelatin-coated plates in IMDM/FBS medium.^71^ For endoderm induction, ESCs or EpiSCs were plated on gelatin-coated plates in either IMDM/FBS or GMEM supplemented with 10% FBS.

### Derivation of Sox1 positive mouse Neural Stem Cells

Differentiation was carried out using Sox1-GFP-puromycin-resistance knock-in mouse (46C) ESC through an adherent monolayer culture in N2B27 medium, consisting of a 1:1 mixture of DMEM/F12 and Neurobasal media, supplemented with 0.5×N2, 0.5×B27, 1% L-glutamin, and 0.1 mM β-mercaptoethanol, as described previously. Under these serum-free conditions, Sox1-GFP positive, puromycin-resistant Rosette NSCs appeared between days 5 and 7. A transient selection with 1.0 μg/ml puromycin, followed by fluorescence-activated cell sorting (FACS) using the BD FACSAria III system, effectively eliminated Sox1-negative cells, resulting in a highly enriched population of Sox1-GFP positive mouse NSCs at Passage 0. These cells were then cultured as neurospheres in Petri dishes with N2B27 medium, supplemented with bFGF/EGF.

### Flow cytometry analysis

ESC (RFP-Zeocin) and CD1-EpiSC (GFP-IP) were dissociated using 0.025% trypsin (GIBCO). Following dissociation. After a single wash with PBS, the cells were resuspended in PBS containing 2% fetal bovine serum with DAPI for analysis. Finally, the stained cells were analyzed using a flow cytometer.

### Immunofluorescence analysis

Cells were fixed on plates with 4% paraformaldehyde (PFA) for 15 minutes at room temperature (RT), followed by three washes with PBS. Blocking was performed using 5% BSA in PBS containing 0.3% Triton X-100 for 1 hour. Primary and secondary antibodies were incubated either for 1 hour at RT or overnight at 4°C, with dilutions prepared in 1% BSA in PBS containing 0.3% Triton X-100. The antibodies used are listed in the Key Resources Table. Cell imaging was conducted using Keyence BZ-X800 fluorescence microscope.

### Alkaline phosphatase (AP) staining

The AP staining reagent was prepared with 200 mM Tris-HCl buffer (pH 8.2) according to the manufacturer’s instructions. Cells were washed with PBS and then incubated in the AP staining reagent (Vector Laboratories, SK-5300) for 30 minutes at room temperature in a darkroom. Following incubation, the cells were rinsed with PBS and fixed with 4% formaldehyde at 4°C overnight. After two PBS washes, the cells were observed using Keyence BZ-X800 fluorescence microscope.

### Single-cell RNA-seq

Co-cultured cells were trypsinized and dissociated into the single-cell suspension in DMEM supplemented with 10% FBS. Dissociated cells were subjected to single cell RNA-seq following the manufacturer’s instructionsusing the 10x Genomics Chromium Next GEM Single Cell 3’ Kit v3.1.

### Cut&Tag

For Cut&Tag, co-cultured ESC- and EpiSC-derived cells were separated based on GFP and RFP fluorescence by FACS sorting. 2x10^5^ cells were used for Cut&Tag for H3K4me3, H3K27me3, H3K27ac using the Epicypher CUTANA CUT&Tag Kit. Sequencing was carried out on Illumina sequencers.

## QUANTIFICATION AND STATISTICAL ANALYSIS

### Single-cell analysis

Single-cell RNA-seq fastqs were aligned to the mm39 genome and processed to barcoded count files using 10X Genomics cellranger 7.2 software. Custom eGFP and RFP sequences were added to the mm39 genome to identify cells with eGFP and RFP expression, respectively. Single-cell count matrices were analyzed using Seurat version 4.3. Prior to SCT transformation, cells with less than 4000 features, less than 10000 counts, and greater than 10% mitochondrial RNA were removed. A total of 4225 cells remained after filtering. Gene expression UMAP plots (Figure 3, panels B and C) were created using the Seurat FeaturePlot function. Naïve cell markers were Nanog, Esrrb, Zfp42, Nr0b1, Tfcp2l1, Tbx3, Tcl1 and Prdm14. Primed cell markers were Fgf5 and Pitx2. Vimentin was used as a MEF cell marker. A UMAP showing the 6 unique cell clusters was created with the Seurat DimPlot function (Figure 3, panel D).

### RNA-seq data analysis

For public RNA-seq data, Raw FASTQ files for public RNA-seq data were downloaded from GEO (GSE45719, GSE100597, GSE74155, and GSE131553). Gene read counts (TPM normalized) were obtained using Salmon v0.14.1 and GENCODE vM24. All data sets were merged, and quantile normalization was performed using the “normalize.quantiles” function from the “preprocessCore” R package. Batch correction was performed using the “removeBatchEffect” function from the “limma” R package. The UMAP plot with the merged data sets was created using the “umap” R package. First, the UMAP base was generated using the “umap” function and samples from GSE45719^73^ and GSE100597^74^. Next, the additional samples were added using the “predict” function. For the correlation plot, genes were first filtered to only include the top 2,000 genes with the highest average expression. Next, the Spearman correlation was calculated using the average expression across cells for each cell type.

For RNA-seq data generated in this study, the raw reads were trimmed with cutadapt (v3.7) first, and the filtered reads were then aligned to mm9 reference with STAR (v2.7.0b). Differentially expressed genes were identified using DESeq2 v1.40.2 and required an FDR of less than 0.05 and a log2 fold change of 1.5 or greater. The expression heatmap was generated using the “heatmap.2” function from the “plots” R package. The average expression was Z-score normalized for each gene. The heatmap only used genes that were differentially expressed in both EpiSC compared to 2iESC and also scPrime compared to scNaive.

### ChIP-seq and Cut&Tag data analysis

For public ChIP-seq data, reads were filtered to only include those with a mean PHRED quality score of 20 or greater. Adapter was trimmed from reads using Cutadapt v4.5. Reads were aligned using Bowtie 2 v2.5.2 and the mm10 reference genome with parameters: “--local -- very-sensitive --no-mixed --no-discordant”. Read coverage was derived from the fully aligned fragments using the “genomecov” tool from the bedtools suite v2.31.1. Read coverage was normalized to depth per 10 million aligned reads. Peaks were called using MACS2 v2.2.9.1. Naïve-specific peaks were defined as peaks with at least 4 times as many depth-normalized reads in the Naïve sample as the Primed sample (and vice versa for Primed-specific peaks). Heatmaps were generated using deepTools v3.5.4.

Cut&Tag analysis was carried out based on instructions from the Epicypher CUTANA CUT&Tag Kit.

### Quantification and Statistical Analysis

All data are presented as mean ± SEM. A two-tailed Student’s t-test was employed to assess statistical significance, with error bars representing the SEM of three independent experiments. A P-value of less than 0.05 was considered statistically significant. Statistical significance is indicated as follows: *P < 0.05, **P < 0.01, and ***P < 0.001.

## ACKNOWLEDGMENTS

We thank the members of the Ying lab for their technical support. This work was supported by the National Institutes of Health (R01 GM129305), the Chen Yong Foundation of the Zhongmei Group, the Xia Research Fund, and the Wu & Jian Research Fund. This study was supported in part by the Intramural Research Program of the NIH, National Institute of Environmental Health Sciences Z01ES102745 (to G.H.).

## AUTHOR CONTRIBUTIONS

Conceptualization, D.W. and Q.-L.Y.; D.W. designed the experiments and performed the majority of the experiments with assistance from X.C., Y.C., J.F., and J.T. X.W. and G.H. performed bulk RNA-seq and single-cell RNA-seq. S.M. performed CUT&Tag. X.W. and G.H. conducted bioinformatic analyses. S.W. and K.S. performed mouse microinjection experiments. Y.C. assisted D.W. with flow cytometry analysis. C.Z., X.C., and D.M. contributed to project discussions. Q.-L.Y. and G.H. provided research support. D.W. and Q.-L.Y. co-wrote the manuscript, with X.W., G.H., and D.M. assisting in manuscript revision.

## Supplemental Figure Legends

**Figure S1.**
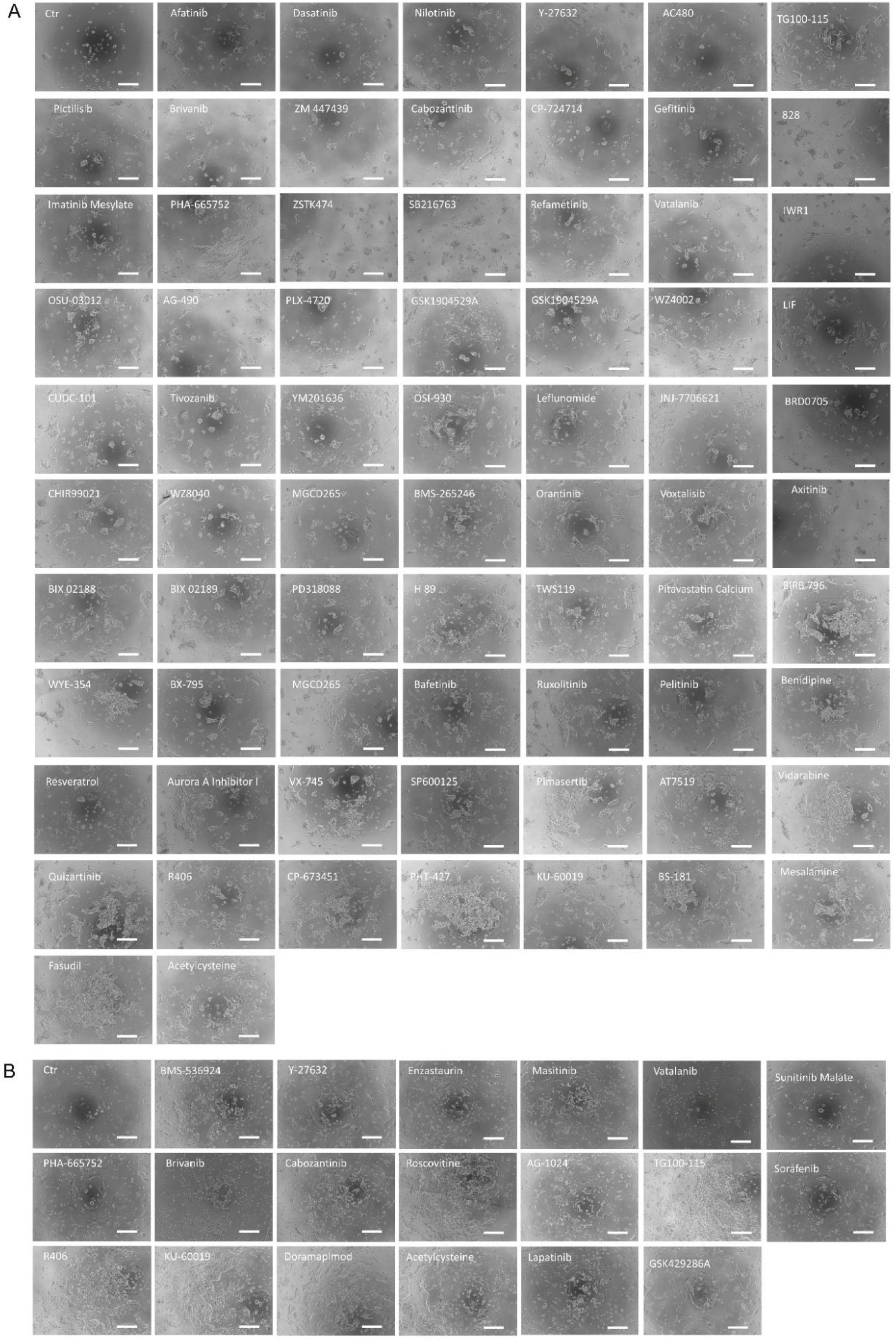
Representative Phenotypes of ESC and EpiSC Treated with Self-Renewal-Promoting Small Molecules. Representative phenotypes of cells treated with small molecules from the library that are effective for the self-renewal of ESCs and EpiSCs. Figure A represents ESCs, and Figure B represents EpiSCs. Scale bars, 250 μm.

**Figure S2.**
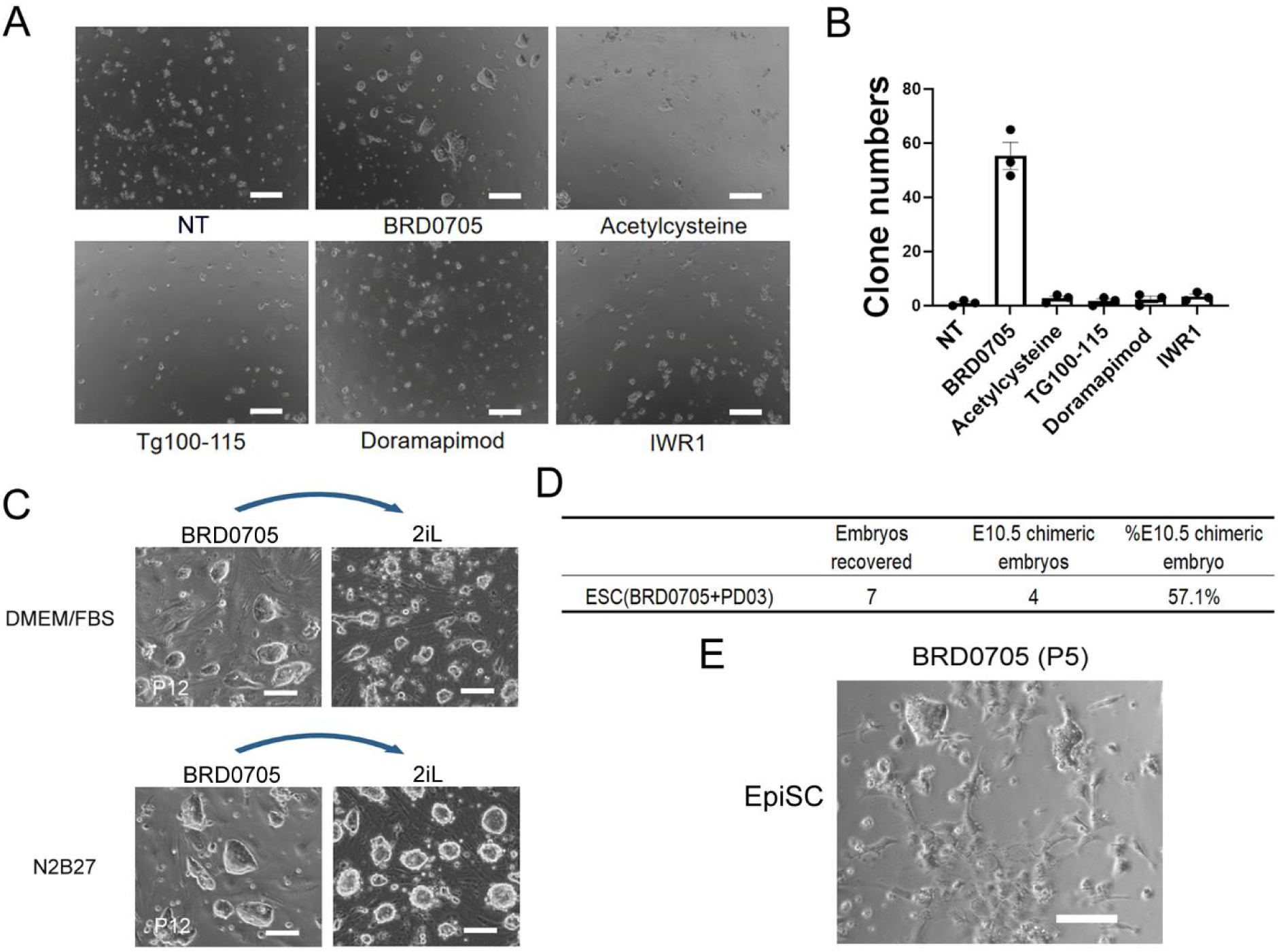
The role of BRD0705 in Maintaining the Self-renewal of ESC and EpiSC. (A) The cellular morphology of ESC treated with the specified compound from the chemical inhibitor library. Cells are at passage 3 under the above conditions, cultured in DMEM/FBS. Scale bars, 200 μm. (B) Quantification of (A). Clone numbers of ESC under different treatments. (C) The left panel shows phase-contrast microscopy images of Passage 12 B6D2F1 ESC (Offspring mice from the cross between BDA male mice and C57BL/6 female mice) cultured in BRD0705 with either N2B27 medium or DMEM/FBS medium. The right panel shows the morphology of ESC after 12 passages in N2B27 and DMEM/FBS media with BRD0705, followed by a change in culture conditions to CHIR/PD03/LIF for three passages. Scale bars, 100 μm. (D) Summary of chimera experiments at E10.5. (E) Representative images showing the morphology of EpiSC after five passages in DMEM/FBS/BRD0705 culture conditions. Scale bars, 100 μm.

**Figure S3.**
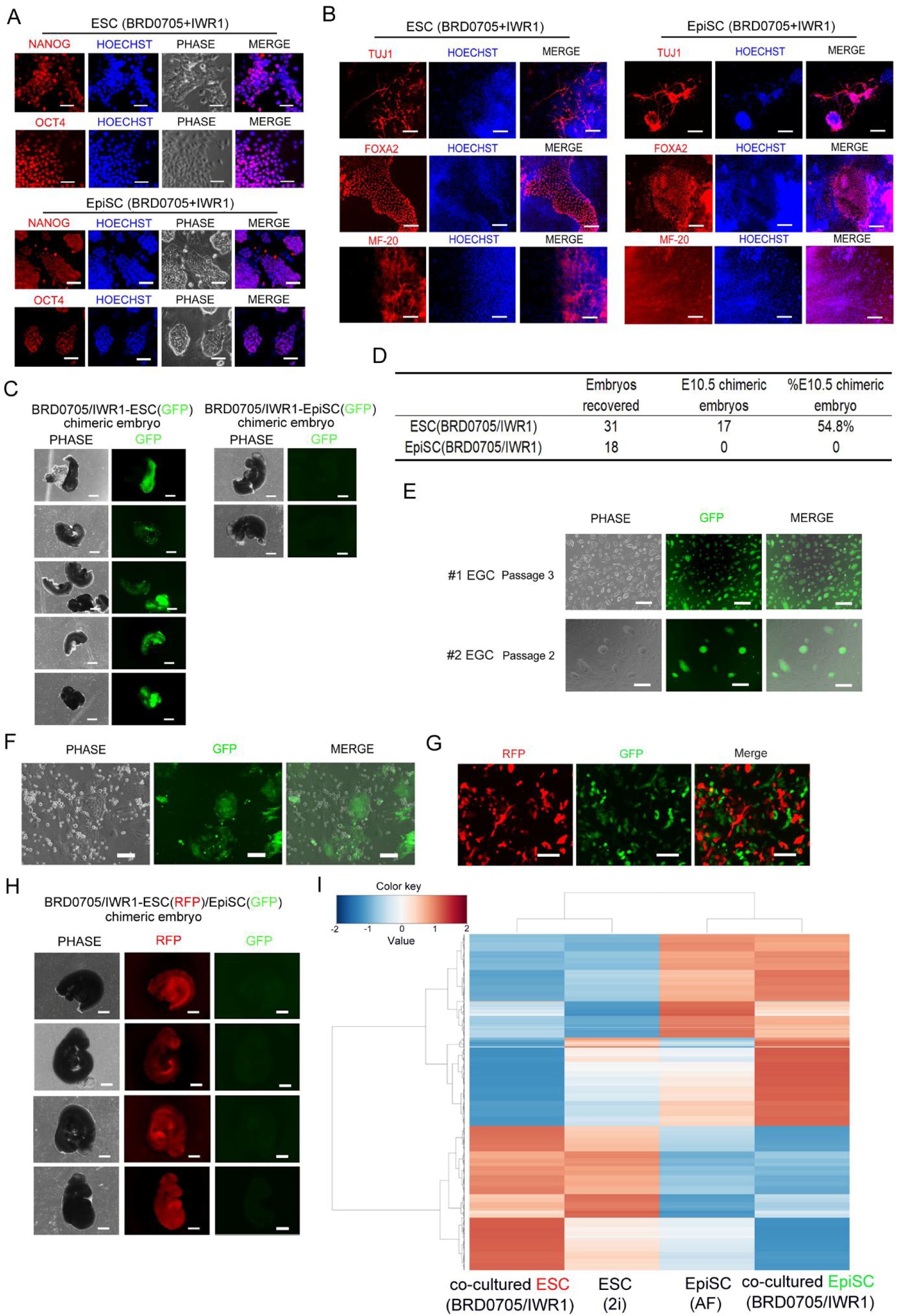
Characterization of ESC and EpiSC Identity under BRD0705/IWR1. (A) IF images confirm the expression of pluripotency markers NANOG and OCT4 in ESCs and EpiSCs cultured with BRD0705/IWR-1 for 12 passages. Scale bar: 50 μm. (B) IF images demonstrate three germ layers differentiation potential of ESC and EpiSC cultured in BRD0705/IWR-1, showing differentiation into endoderm (FOXA2), mesoderm (MF-20), and ectoderm (TUJ1). Scale bar: 250 μm (C) Representative phase-contrast and fluorescence images of E10.5 mouse embryos derived from WT blastocyst injection of GFP-labeled BRD0705/IWR1-ESC or GFP-labeled BRD0705/IWR1-EpiSC. Scale bars, 500 μm. (D) Summary of chimera experiments at E10.5. (E) Phase-contrast and fluorescence images of EGCs derived from two different gonads (gonads obtained from two E15.5 chimeric mouse embryos produced via wild-type blastocyst injection of GFP-labeled ESC cultured with BRD0705/IWR1), with cells cultured under 2i/LIF conditions. Scale bars, 250 μm. #1 and #2 represent EGCs derived from the gonads of two distinct chimeric mice. (F) Representative phase-contrast and fluorescence images of mixed cells derived from E10.5 mouse embryos obtained from WT blastocyst injections of GFP-expressing ESC cultured with BRD0705/IWR1. The image shows beating GFP-positive muscle cells, with a video of the beating cells provided in Supplementary Movie S1. Scale bars, 100 μm (G) The phase-contrast and fluorescence images of Passage 1 ESC (RFP+)/EpiSC (GFP+) cultured in N2B27 medium supplemented with BRD0705/IWR-1, respectively. Scale bars, 100 μm. (H) Representative images of phase-contrast and fluorescence microscopy of E10.5 mouse embryos following blastocyst injection with a 1:1 mixture of RFP-labeled ESC and GFP-labeled EpiSC, cultured in the presence of BRD0705/IWR1. Scale bars, 500 μm. (I) Heatmap showing the transcriptomic correlation of ESC and EpiSC co-cultured under BRD0705/IWR1 conditions compared to reference ESC (2i) and EpiSC (AF). The gene expression profiles of co-cultured ESCs and EpiSCs closely resemble those of ESCs cultured in 2i and EpiSCs cultured in AFX, respectively, indicating that BRD0705/IWR1 maintains the transcriptional identity of both cell types.

**Figure S4.**
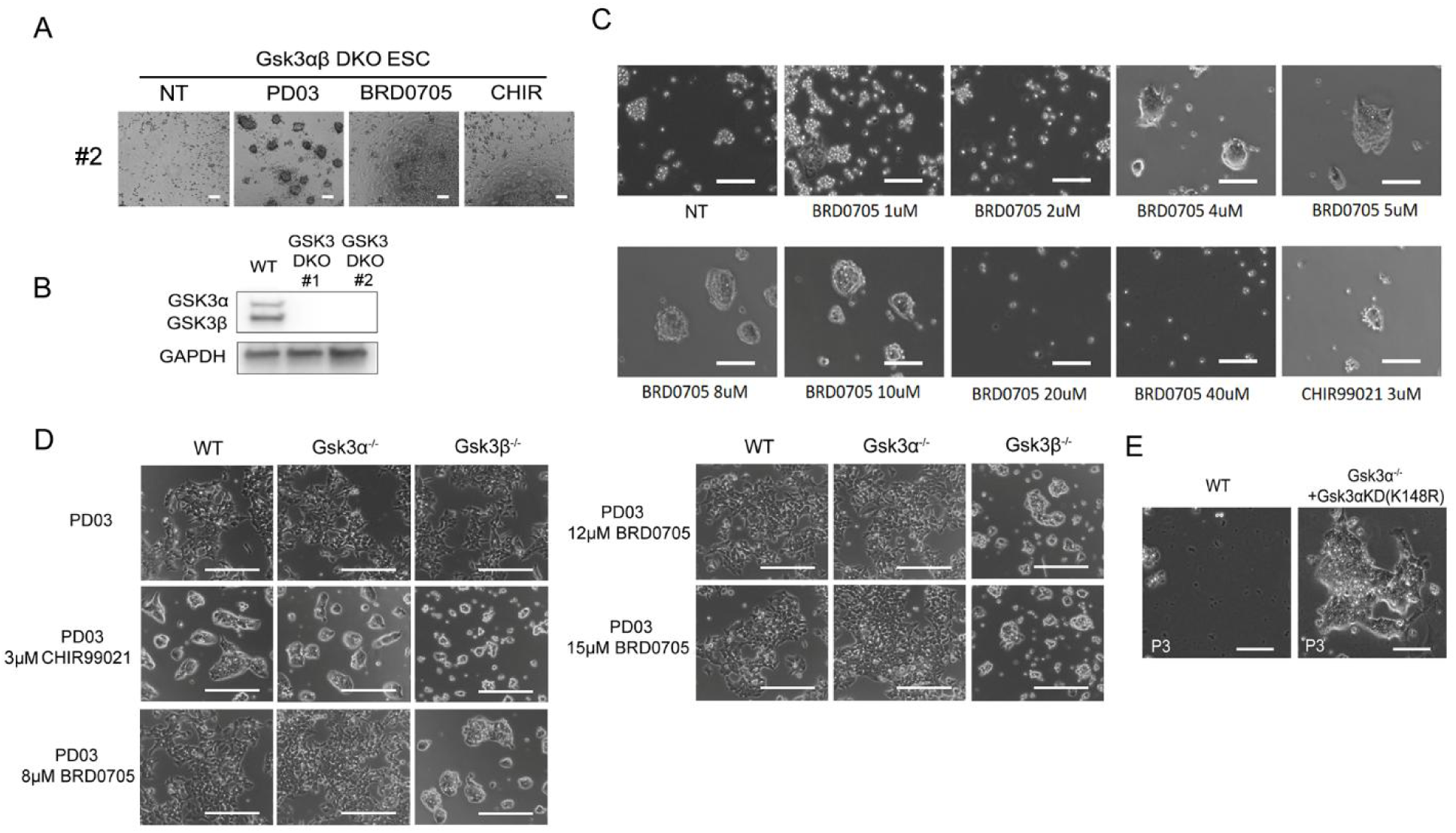
BRD0705 Regulates ESC Self-Renewal Through Gsk3α Inhibition. (A) Morphological comparison of *Gsk3α/β* DKO ESC clones under different culture conditions. Representative images of #2 Gsk3αβ DKO ESC line cultured in DMEM/FBS medium with NT, PD03, BRD0705, and CHIR treatments for 3 passages. Scale bars, 100 μm. (B) Western blot analysis of GSK3α and GSK3β in wild-type (WT) and *GSK3α/β* DKO cell line #1 and #2. GSK3α and GSK3β are present in WT but absent in DKO samples, confirming the knockout. GAPDH serves as control. (C) Dose-dependent effects of BRD0705 on Gsk3β^−/−^ ESC morphology. Representative images show Gsk3β^−/−^ ESCs cultured in N2B27/PD03 with BRD0705 (1–40 μM) for 2 passages, compared to CHIR99021 (3 μM). Scale bars, 100 μm. (D) Morphological analysis of WT, Gsk3α^−/−^, and Gsk3β^−/−^ ESC cultured in N2B27 medium with PD03, or combined with CHIR99021 or BRD0705 at 8 µM, 12 µM, and 15 µM for 3 passages. Representative images display differences in colony morphology and cell density. Scale bars: 200 µm. (E) Morphology of wild-type (WT) and Gsk3α^−/−^ cells with kinase-dead Gsk3α (K148R) mutation at passage 3 (P3). Cell cultured in DMEM/FBS medium for 3 passages. After 3 passages, all WT ESC have died, whereas Gsk3α^−/−^+Gsk3αKD cells still maintain the presence of colonies. Scale bars represent 100 µm.

**Figure S5.**
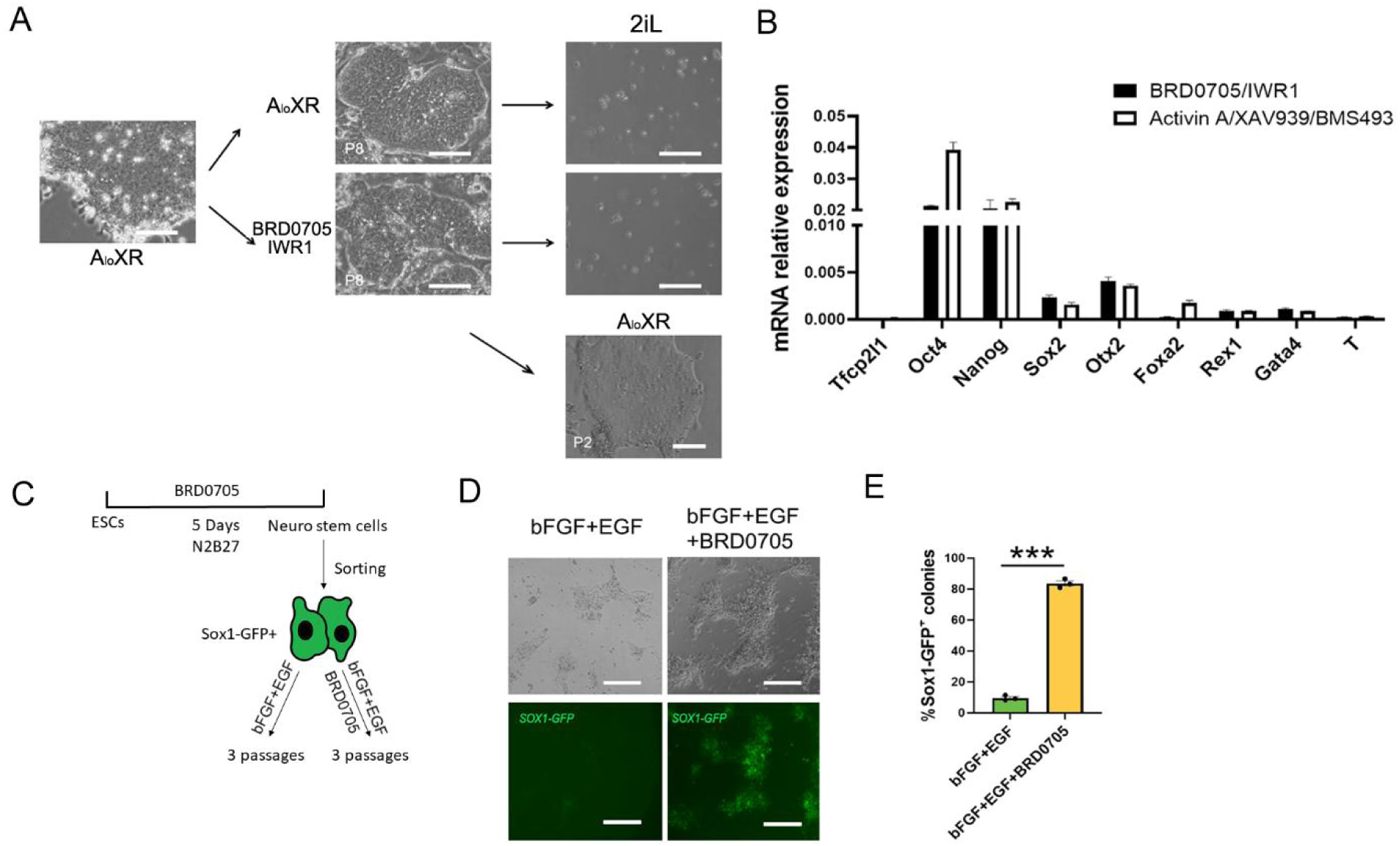
BRD0705 Effectively Maintains the Stemness of Formative cell and Neural Stem Cells. (A) Representative images showing the morphology of formative cells cultured under BRD0705/IWR-1 conditions or maintained in A_lo_XR conditions for eight passages. They were then transitioned to 2iL conditions to assess cell viability, while cells cultured under BRD0705/IWR1 were also reverted to A_lo_XR as a control. Scale bars, 100 μm. (B) qRT-PCR analysis of marker gene expression was performed on formative cells cultured under both BRD0705/IWR-1 and A_lo_XR conditions, with expression levels normalized to GAPDH. Error bars represent the SEM from technical triplicates. (C) Schematic illustration of the experimental design. SOX1-GFP+ neural stem cells were isolated from 46C ESCs on day 5 of differentiation and then expanded under bFGF+EGF conditions, either with or without BRD0705. (D) Representative images illustrating the impact of BRD0705 treatment on neural stem cell expansion. Scale bars, 200 μm. (E) Quantification of Figure O. The bar graph shows the percentage of Sox1-GFP+ colonies in neural stem cells under control (Ctr) and BRD0705 treatment conditions (*, p < 0.05). Data are presented as mean ± SEM.

**Table S1.** List of small-molecule library compounds.

**Table S2.** List of qPCR primers.

**Movie S1.** Beating of GFP+ muscle cells in E10.5 mouse embryos obtained by injecting wild-type blastocysts with GFP-labeled ESCs cultured with BRD0705/IWR1.

